# Increased RNA and protein degradation is required for counteracting transcriptional burden and proteotoxic stress in human aneuploid cells

**DOI:** 10.1101/2023.01.27.525826

**Authors:** Marica Rosaria Ippolito, Johanna Zerbib, Yonatan Eliezer, Eli Reuveni, Sonia Viganò, Giuseppina De Feudis, Anouk Savir Kadmon, Ilaria Vigorito, Sara Martin, Kathrin Laue, Yael Cohen-Sharir, Simone Scorzoni, Francisca Vazquez, Stefano Santaguida, Uri Ben-David

## Abstract

Aneuploidy, an abnormal chromosome composition, results in a stoichiometric imbalance of protein complexes, which jeopardizes the fitness of aneuploid cells. Aneuploid cells thus need to compensate for the imbalanced DNA levels by regulating their RNA and protein levels, a phenomenon known as dosage compensation. However, the molecular mechanisms involved in dosage compensation in human cells – and whether they can be targeted to selectively kill aneuploid cancer cells – remain unknown. Here, we addressed this question via molecular dissection of multiple diploid vs. aneuploid cell models. Using genomic and functional profiling of a novel isogenic system of RPE1-hTERT cells with various degrees of aneuploidy, we found that aneuploid cells cope with both transcriptional burden and proteotoxic stress. At the mRNA level, aneuploid cells increased RNA synthesis, but concomitantly elevated several RNA degradation pathways, in particular the nonsense-mediated decay (NMD) and the microRNA-mediated mRNA silencing pathways. Consequently, aneuploid cells were more sensitive to the genetic or chemical perturbation of several key components of these RNA degradation pathways. At the protein level, aneuploid cells experienced proteotoxic stress, resulting in reduced translation and increased protein degradation, rendering them more sensitive to proteasome inhibition. These findings were recapitulated across hundreds of human cancer cell lines and primary tumors, confirming that both non-transformed and transformed cells alter their RNA and protein metabolism in order to adapt to the aneuploid state. Our results reveal that aneuploid cells are dependent on the over- or under-activation of several nodes along the gene expression process, identifying these pathways as clinically-actionable vulnerabilities of aneuploid cells.

## Introduction

Aneuploidy is a genomic state characterized by chromosome gains and losses. A major consequence of aneuploidy is genome and proteome imbalance, which aneuploid cells must overcome in order to function properly. The degree of gene dosage compensation varies across different cellular contexts^1^, yet it is clear that in human aneuploid cancer cells the effect of aneuploidy is attenuated by such buffering mechanisms. Recent studies have revealed that many proteins do not change their expression by the degree expected based on their DNA levels^2–6^. The mechanisms that allow for dosage compensation, and the potential cellular vulnerabilities that result from them, remain under-explored.

Previous studies have exposed the role of protein regulation and protein degradation for “buffering” the effect of copy number alterations (CNAs). Aneuploid cells experience proteotoxic stress, which is partly overcome in aneuploid yeast by an increased activity of the proteasome^7–10^. Similarly, a recent study described a protein folding deficiency in engineered aneuploid human cells^2^. However, the role of the proteasome in the context of aneuploid human cancer cells has remained unknown, and is of particular clinical relevance given that proteasome inhibitors are used in the clinic (mostly for treating multiple myeloma)^11^. It also remains unknown whether other important processes of protein metabolism, such as protein translation, are also dysregulated in aneuploid cells.

Gene expression is, of course, also regulated at earlier stages of mRNA regulation. Whereas dosage compensation at the mRNA level seems to be minimal in yeast^7,12,13^, it does occur in human cancer cells^4,5,14,15^ : a recent analysis of cancer cell lines found that the mRNA expression of ∼20% of the genes does not scale with their chromosome-arm copy number levels^4^, and a similar percentage of such genes was observed in human primary tumor data^14^. However, the potential role of RNA transcription, metabolism and degradation in attenuating aneuploidy-induced gene expression changes – and whether this can create cellular vulnerabilities in aneuploid cells – have yet to be explored.

In our companion study, we established a library of stable RPE1 clones with various degrees of aneuploidy (Zerbib, Ippolito et al, *bioRxiv* 2023). Here, we analyzed genomic and functional data from these isogenic clones and uncovered an increased vulnerability of aneuploid cells to perturbation of RNA and protein degradation pathways. These novel aneuploidy-induced functional dependencies were validated in human cancer cell lines, and differential activity of these pathways was confirmed in primary human tumors. These findings may thus have important clinical ramifications, both for the development of novel cancer therapeutics and for predicting patients’ response to existing drugs.

## Results

### Increased RNA synthesis and degradation in aneuploid cells

To investigate dosage compensation in aneuploid cells, we used a novel isogenic system of non-transformed chromosomally stable aneuploid cells, presented in detail in our companion study (Zerbib, Ippolito et al, *bioRxiv* 2023). Briefly, we transiently treated RPE1-hTERT cells with the MPS1 inhibitor reversine to induce chromosome mis-segregation and generate aneuploidy^16–18^, single-cell sorted and karyotyped the obtained clones (**Fig. 1a** and (Zerbib, Ippolito et al, *bioRxiv* 2023). We selected 7 clones with increasing degrees of aneuploidy: three pseudo-diploid clones, RPE1-SS48, RPE1-SS31 and RPE1-SS77 (hereinafter SS48, SS31 and SS77, respectively), two clones carrying a single extra chromosome, RPE1-SS6 and RPE1-SS119 (hereinafter SS6 and SS119, respectively), and two clones carrying multiple trisomies, RPE1-SS51 and RPE1-SS111 (hereinafter SS51 and SS111, respectively). We characterized the clones extensively, demonstrating their high relevance for aneuploidy research (Zerbib, Ippolito et al, *bioRxiv* 2023).

**Figure 1:**
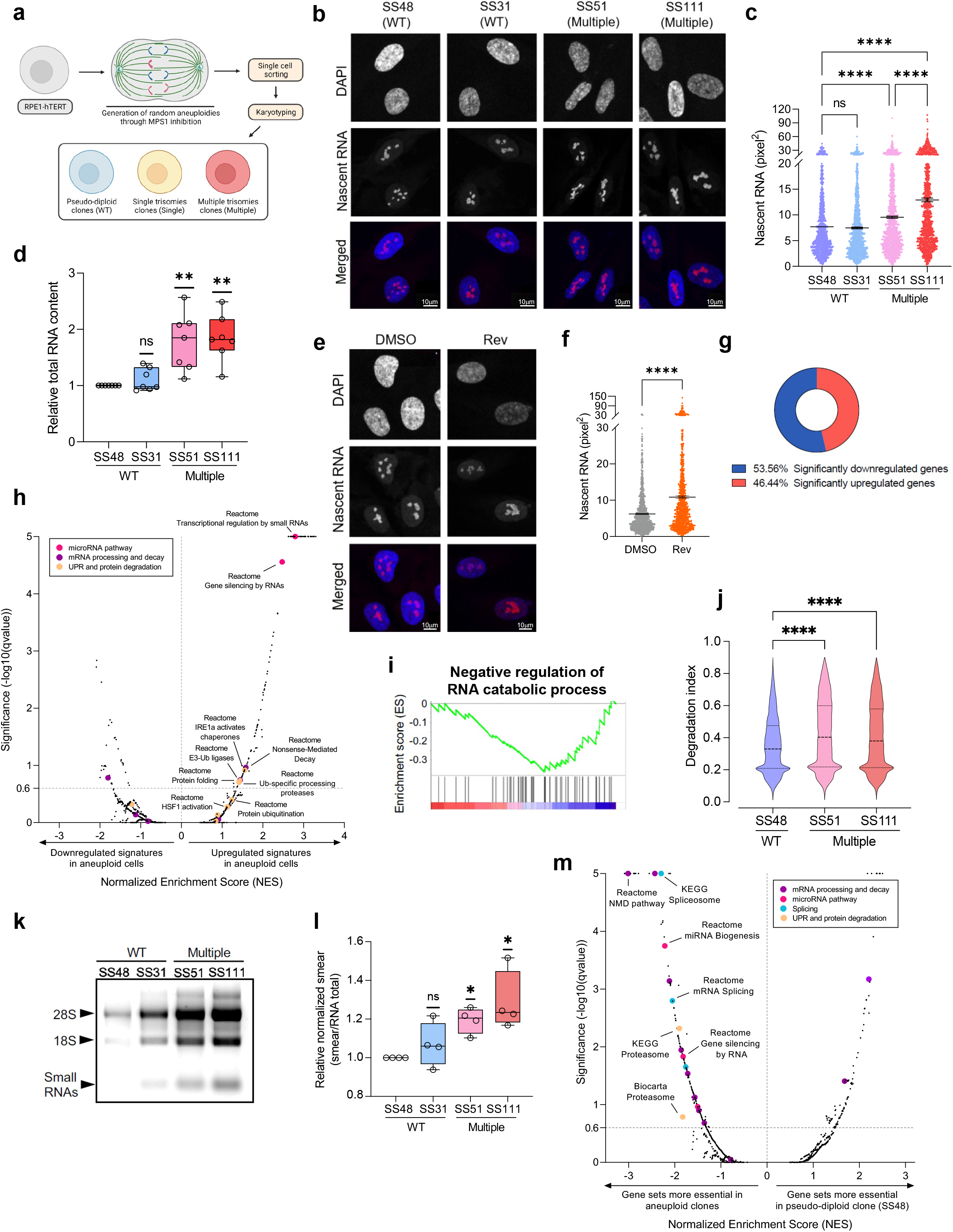
Aneuploid cells compensate for the extra DNA content through increased RNA and protein turn-over. **(a)** Schematic representation of clone generation. See Zerbib, Ippolito et al. *bioRxiv* 2023 for more details. **(b)** Immunofluorescence of nascent RNA foci in pseudo-diploid clones, SS48 and SS31, and in highly-aneuploid clones, SS51 and SS111. Red, nascent RNA; Blue, DAPI; Scale bar, 10um. **(c)** Quantitative comparison of nascent RNA showing area (pixel) of nascent RNA foci. n=3 independent experiments; ****, p<0.0001; Kruskal–Wallis test, Dunn’s multiple comparison. **(d)** Quantification of total RNA between pseudo-diploid clones (SS48 and SS31) and highly-aneuploid clones (SS51 and SS111). n= 7 independent experiments; RNA content was calculated relative to SS48, per experiment. **, p=0.007 and p=0.0018, for SS51 and SS111 respectively; One-Sample t-test. **(e)** Immunofluorescence of nascent RNA foci in pseudo-diploid RPE1-hTERT treated with DMSO or after 72hrs following reversine pulse. Red, nascent RNA; Blue, DAPI; Scale bar, 10um. **(f)** Quantitative comparison of nascent RNA showing area (pixel) of nascent RNA foci. n=3 independent experiments; ****, p<0.0001; two-tailed Mann-Whitney test. **(g)** The fractions of significantly upregulated and downregulated genes out of all differentially-expressed genes, between the highly-aneuploid clones, SS51 and SS111, and the pseudo-diploid clone SS48. n=8215 genes (qvalue<0.25). ****, p<0.0001; two-tailed binomial test. Data are obtained from Zerbib, Ippolito et al, *bioRxiv* 2023. **(h)** Comparison of the differential gene expression patterns (pre-ranked GSEA results) between the pseudo-diploid SS48 clone (control) and the highly-aneuploid SS51 and SS111 clones. Plot presents enrichments for the Hallmark, KEGG, Biocarta and Reactome gene sets. Data are obtained from Zerbib, Ippolito et al, *bioRxiv* 2023. Significance threshold set at qvalue=0.25. Enriched pathways are color-coded. **(i)** Gene set enrichment analysis (GSEA) of an RNA catabolism gene expression signature, comparing the highly-aneuploid clones, SS51 and SS111, to the pseudo-diploid clone SS48. Data are obtained from Zerbib, Ippolito et al, *bioRxiv* 2023. Shown is an enrichment plot for the GO Biological Process ‘Negative regulation of RNA catabolic processes’ gene set (NES= -1.58; q-value=0.2). **(j)** Comparison of the mean degradation index (degraded RNA score) across all genes (n=13,689), using the Degnorm algorithm. ****, p<0.0001; Repeated-Measured One-Way ANOVA, Tukey’s multiple comparison test. **(k)** Native agarose gel electrophoresis of total RNA extracted from RPE1 clones, re-suspended in Nuclease-Free water, showing a specific increased amount of RNA smear in the highly-aneuploid clones, SS51 and SS111, in comparison to the pseudo-diploid clones SS48 and SS31. **(l)** Quantification of RNA degradation, as evaluated by the smear/total RNA ratio. Fold change in normalized smear was calculated relative to SS48, per experiment. n=4 independent experiments; *, p=0.0102 and p=0.034, for SS51 and SS111, respectively; One-Sample t-test. **(m)** Comparison of the differential gene dependency scores (pre-ranked GSEA results) between the near-diploid SS48 clone (control) and the aneuploid SS6, SS119 and SS51 clones. Plot presents enrichments for the Hallmark, KEGG, Biocarta and Reactome gene sets. Data are obtained from Zerbib, Ippolito et al, *bioRxiv* 2023. Significance threshold set at qvalue=0.25. Enriched pathways are color-coded.

As the selected aneuploid clones carry extra chromosomes, we hypothesized that this excessive DNA content would lead to increased RNA synthesis in these cells. We focused on the most aneuploid clones, SS51 (trisomic for chromosomes 7 and 22) and SS111 (trisomic for chromosomes 8, 9 and 18), and quantified newly synthesized RNA in our models using Ethynyl Uridine (5-EU) incorporation. Indeed, nascent RNA was more abundant in highly aneuploid clones, with the highest synthesis levels found in the most aneuploid clone, SS111 (**Fig. 1b-c**). In line with these findings, the total levels of extracted RNA were higher in the highly-aneuploid clones in comparison to pseudo-diploid clones (**Fig. 1d**), consistent with previous studies showing the correlation between DNA and RNA content in aneuploid cells^4,19,20^. To assess whether increased RNA synthesis is an immediate consequence of aneuploidy, we quantified the newly synthesized RNA in parental RPE1-hTERT cells (hereinafter parental RPE1 cells) 72hrs following a pulse of reversine. Interestingly, reversine-treated RPE1 cells also increased their nascent RNA levels (**Fig. 1e-f**), in agreement with the results obtained in the stable aneuploid clones.

Next, we investigated the gene expression differences between the pseudo-diploid and aneuploid RPE1 clones, using genome-wide RNA sequencing (RNAseq). Despite their increased transcription, differential gene data analysis revealed that more genes were downregulated than upregulated in the highly-aneuploid clones, independently of p53 mutation status (p<0.001; **Fig. 1g** and **Extended Data Fig. 1a**), suggesting gene dosage compensation at the mRNA level in our model. As our aneuploid RPE1 clones harbor different trisomies, we then applied pre-ranked gene set enrichment analysis (GSEA^21,22^) to identify gene expression signatures that are induced by aneuploidy regardless of the specific affected chromosome(s). The most elevated transcriptional signatures in the aneuploid clones were associated with RNA and protein regulation. Specifically, we identified a significant upregulation of signatures related to RNA metabolism and gene silencing, *e*.*g*. ‘nonsense mediated decay’ and ‘gene silencing by RNAs’ (**Fig. 1h, Extended Data Fig. 1b**), and to the unfolded protein response (UPR) and protein degradation, e.g. ‘IRE1a activates chaperones’ and ‘E3-Ub ligases ubiquitinate target proteins’ (**Fig. 1h, Extended Data Fig. 1b**). These results suggest global attenuation of gene and protein expression in the aneuploid clones, consistent with previous studies^3,4,20,23,24^.

Thus, we investigated RNA degradation in the pseudo-diploid vs. highly-aneuploid clones. Gene set enrichment analysis showed increased RNA catabolism in highly-aneuploid cells in comparison to their pseudo-diploid counterparts (**Fig. 1i**). We therefore leveraged our global RNAseq data to quantify RNA degradation in the samples using ‘DegNorm’, an algorithm developed to quantify degraded RNA and remove its effect from RNAseq data analyses^25^. We found a significant increase in the RNA degradation index (a measure for RNA degradation levels) in the highly-aneuploid clones (**Fig. 1j**). We validated this finding by running a gel electrophoresis on the total RNA extracted from the clones and quantifying the resultant ‘smears’ (**Fig. 1k-l**). We note that RNA degradation levels were highest in the most aneuploid clone, SS111, which also exhibited the highest levels of RNA synthesis (**Fig. 1b-c**). These findings indicate that the increased DNA content in the aneuploid clones with extra chromosomes leads to elevated levels of both RNA synthesis and RNA degradation, resulting in higher RNA turnover in these cells.

Importantly, to confirm that the enrichments found in our RNAseq data analysis were not confounded by the increased levels of RNA degradation in the aneuploid clones, we repeated all differential gene expression analyses after computationally removing the degraded transcripts. We were able to recapitulate the enrichments for DNA damage response (Zerbib, Ippolito et al, *bioRxiv* 2023), RNA metabolism and protein degradation signatures (**Extended Data Fig. 2**).

Finally, we set out to identify genes that are preferentially essential in aneuploid cells, using genome-wide CRISPR/Cas9 screens of the isogenic RPE1 clones (Zerbib, Ippolito et al, *bioRxiv* 2023). Consistent with their gene expression profiles, unbiased pre-ranked GSEA analysis revealed that aneuploid clones were more dependent on genes related to gene silencing through RNA processing and decay, including the nonsense-mediated decay (NMD) pathway, the miRNA pathway, and gene splicing (**Fig. 1m**). Indeed, the increased levels of DNA damage that we identified in the aneuploid clones (Zerbib, Ippolito et al, *bioRxiv* 2023), coupled to increased rate of RNA synthesis, might result in an excessive number of abnormal transcripts, potentially explaining why aneuploid cells would be more dependent on RNA processing and degradation. Moreover, aneuploid clones were also more dependent on protein degradation via the proteasome (**Fig. 1m**), consistent with ongoing proteotoxic stress and the resultant accumulation of aberrant proteins (**Fig. 1h**). Together, these results suggest that cells with extra chromosomes strongly rely on the downregulation of their gene expression to compensate for their extra DNA content, both at the RNA and at the protein level.

### Increased NMD activity and dependency in aneuploid cells

Next, we assessed potential mechanisms of RNA degradation. The highly aneuploid clones, SS51 and SS111, exhibited elevated transcriptional signatures of the NMD pathway (**Fig. 1h, Fig. 2a** and **Extended Data Fig. 3a**). Thus, we compared their NMD pathway activity to that of the pseudo-diploid clones. First, we estimated NMD activity by calculating a transcriptional signature score of described NMD targets^26^. We found a significant increase in this transcriptional score in the highly-aneuploid clones (**Fig. 2b**), consistent with the gene set enrichment analysis (**Fig. 2a**). Next, we validated this increased activity using an NMD pathway reporter system^27^, which confirmed that under standard culture conditions highly-aneuploid clones elevated their NMD pathway activity in comparison to their pseudo-diploid counterparts (**Extended Data Fig. 3b**).

**Figure 2:**
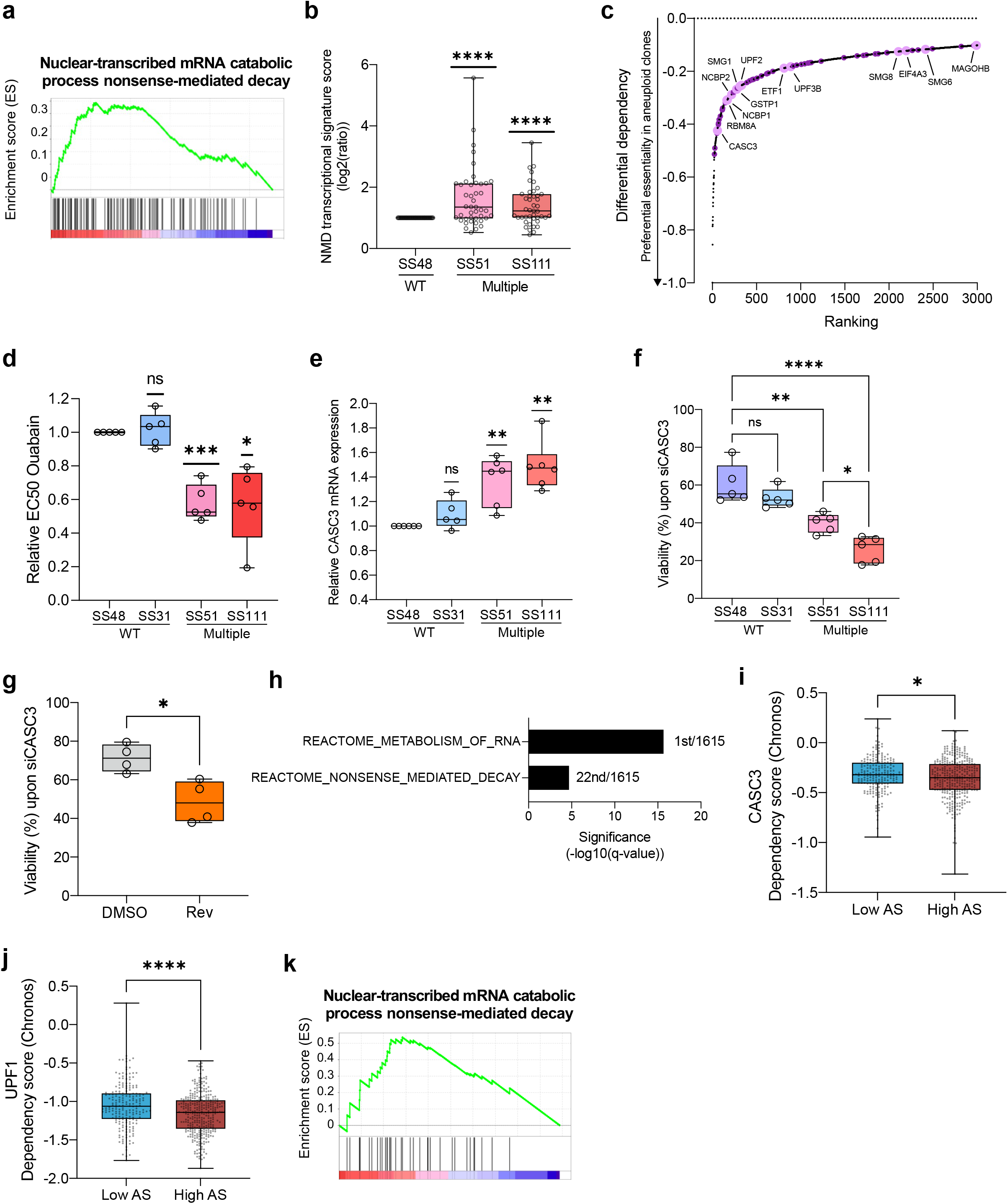
Aneuploid cells activate the nonsense-mediated decay (NMD) pathway, and depend on this pathway for downregulating their gene expression. **(a)** Gene set enrichment analysis (GSEA) of an NMD-related signature, comparing the highly-aneuploid clones, SS51 and SS111, to the pseudo-diploid clone SS48. Shown is the enrichment plot for the GO-Biological Process ‘Nuclear transcribed mRNA catabolic processes NMD’ gene set (NES=1.83; q-value=0.07). Data are taken from Zerbib, Ippolito et al, *bioRxiv* 2023 **(b)** Comparison of gene expression of the NMD pathway between the highly-aneuploid clones SS51 and SS111, and the pseudo-diploid clone SS48. Fold change in transcriptional score was calculated relative to SS48, for each gene (n=43 genes). ****, p<0.0001; One-Sample t-test. Data are obtained from Zerbib, Ippolito et al, *bioRxiv* 2023 **(c)** The top 3,000 genes that aneuploid clones were most preferentially sensitive to their knockout in comparison to the pseudo-diploid clone SS48, based on our genome-wide CRISPR/Cas9 screen. Highlighted are genes that belong to the NMD pathway: core member genes (in pink) and ribosomal-related genes (in purple). NMD-related genes are significantly enriched within the top 3,000 gene list; ****, p<0.0001; two-tailed Fisher’s Exact test. Data are obtained from Zerbib, Ippolito et al, *bioRxiv* 2023. **(d)** Comparison of drug sensitivity (determined by IC50 values) to 72hr drug treatment with the NMD inhibitor ouabain, between pseudo-diploid clones (SS48 and SS31) and highly-aneuploid clones (SS51 and SS111). IC50 fold-change was calculated relative to SS48, per experiment. n=5 independent experiments; *, p=0.012 and ***, p=0.0004, for SS111 and SS51, respectively; One-Sample t-test. **(e)** Comparison of CASC3 mRNA levels, quantified by qRT-PCR, between pseudo-diploid clones (SS48 and SS31) and highly-aneuploid clones (SS51 and SS111). Fold change in CASC3 expression was calculated relative to SS48, per experiment. n=5 (SS31) and n=6 (SS48, SS51, SS111) independent experiments; **, p=0.0058 and p=0.0018, for SS51 and SS111, respectively; One-Sample t-test. **(f)** Comparison of cell viability following siRNA against CASC3 for 72hrs, between pseudo-diploid clones (SS48 and SS31) and highly aneuploid clones (SS51 and SS111). Viability was calculated relative to a control siRNA. n=5 independent experiments; *, p<0.05; **, p<0.01; ****, p<0.0001; One-Way ANOVA, Tukey’s multiple comparison. All comparisons between SS31 and aneuploid clones were significant as well (*, p<0.05). **(g)** Comparison of cell viability following siRNA against CASC3, between parental RPE1 cells treated for 20hrs with the SAC inhibitor reversine (500nM) or with control DMSO, then harvested 72hrs post wash-out. Relative viability was calculated relative to a control siRNA treatment. n=4 independent experiments; *, p=0.0134; two-tailed unpaired t-test. **(h)** Gene set enrichment analysis of the genes whose expression correlates with proliferation in highly-aneuploid cancer cell lines but not in near-diploid cancer cell lines, reveals significant enrichment of multiple RNA metabolism signatures. Shown here are the Reactome ‘Metabolism of RNA’ and ‘Nonsense Mediated Decay’ gene sets. Significance values represent the FDR q-values. The ranking of each RNA metabolism signature (out of all signatures included in the gene set collection) is indicated next to each bar. **(i-j)** Comparison of gene dependency (determined by Chronos score) for key members of the NMD pathway, the EJC member CASC3 (**i**) and the main effector UPF1 (**j**), between the top and bottom aneuploidy quartiles of human cancer cell lines (n=538 cell lines). Data were obtained from DepMap CRISPR screen, 22Q1 release. *, p=0.0289 and ****, p<0.0001, for CASC3 and UPF1 respectively; two-tailed Mann-Whitney test. (**k**) Pre-ranked GSEA of mRNA expression levels showing that high aneuploidy levels are associated with upregulation of the nonsense-mediated decay (NMD) in human primary tumors. Shown is the GO-Biological Process ‘Nuclear transcribed mRNA catabolic processes NMD’ gene set (NES=1.70, q-value=0.029) gene set. Data were obtained from the TCGA mRNA expression data set^62^.

We then turned to investigate the dependency of aneuploid cells on the NMD pathway. The NMD pathway was among the very top differential dependencies of aneuploid cells in the CRISPR screen (**Fig. 1m**), with many of its components ranking among the most differentially-essential genes (**Fig. 2c**). Importantly, these results held true even when the p53-mutated SS77 clone was included in the analysis (**Extended Data Fig. 1c** and **Extended Data Fig. 3c**), indicating that the increased dependency of aneuploid cells on NMD is not simply due to p53 activation. To validate this dependency, we exposed the RPE1 clones to pharmacological inhibitors of NMD, ouabain and digoxin^27^, and found that the highly-aneuploid clones SS51 and SS111 were significantly more sensitive to both drugs (**Fig. 2d** and **Extended Data Fig. 3d**). We then investigated CASC3 (also known as MLN51), the top differentially-essential core member of the NMD pathway, and a key regulator of NMD pathway activation^28^. We found that highly-aneuploid clones upregulated their CASC3 expression in comparison to their pseudo-diploid counterparts (**Fig. 2e**). Moreover, CASC3 protein expression levels increased following reversine-mediated aneuploidization of the parental RPE1 cells, and this increase was observed also in *TP53*-KD and *TP53*-KO RPE1 cells, indicating a p53-independent mechanism (**Extended Data Fig. 3e-h**). Highly-aneuploid clones were significantly more sensitive to genetic CASC3 inhibition by siRNA (**Fig. 2f** and **Extended Data Fig. 3i**). In addition, reversine-induced aneuploidization of the parental pseudo-diploid RPE1 cells also rendered the cells more sensitive to CASC3 inhibition (**Fig. 2g** and **Extended Data Fig. 3j**). Finally, intrigued by previous observations showing that NMD could get activated by the DDR^29–32^, we found that DNA damage induction using etoposide increased CASC3 expression levels in parental RPE1 cells (**Extended Data Fig. 3k-l**), providing a plausible mechanistic link between the increased DNA damage observed in the aneuploid cells (Zerbib, Ippolito et al *bioRxiv* 2023) and their increased expression of, and dependency on, the NMD pathway. Together, these results confirm that aneuploidy increases cellular dependency on the NMD pathway.

Lastly, we asked whether NMD activity and dependency are associated with a high degree of aneuploidy in human cancer cells as well. Gene expression analysis of hundreds of human cancer cell lines revealed that RNA metabolism, and particularly RNA degradation through the NMD pathway, were strongly associated with the proliferation capacity of highly-aneuploid cancer cell lines (but not with that of near-euploid cancer cell lines) (see **Methods, Fig. 2h**). Moreover, analysis of CRISPR screens revealed that highly-aneuploid cancer cells were significantly more dependent on multiple members of the NMD pathway^33^, including CASC3 and the core NMD effector UPF1 (**Fig. 2i-j** and **Extended Data Fig. 3m-p**). Finally, we found a significant association between aneuploidy levels and the NMD signature across human primary tumors as well (**Fig. 2k**). We conclude that NMD activity and dependency are associated with a high degree of aneuploidy in cancer cells.

### Increased miRNA-mediated RNA degradation and altered gene splicing in aneuploid cells

The NMD pathway was not the only RNA degradation pathway that came up in our unbiased genomic and functional analyses. Gene set enrichment analysis also revealed significant enrichment for signatures associated with gene expression silencing through the small RNA pathways (**Fig. 1h** and **Fig. 3a**). Similar to the NMD pathway, the miRNA pathway was among the top differentially-essential pathways in aneuploid cells (**Fig. 1m** and **Fig. 3b**), with the hallmark miRNA pathway genes XPO5, DICER1 and DROSHA scoring among the 20 most differentially-essential genes overall (**Fig. 3b**).

**Figure 3:**
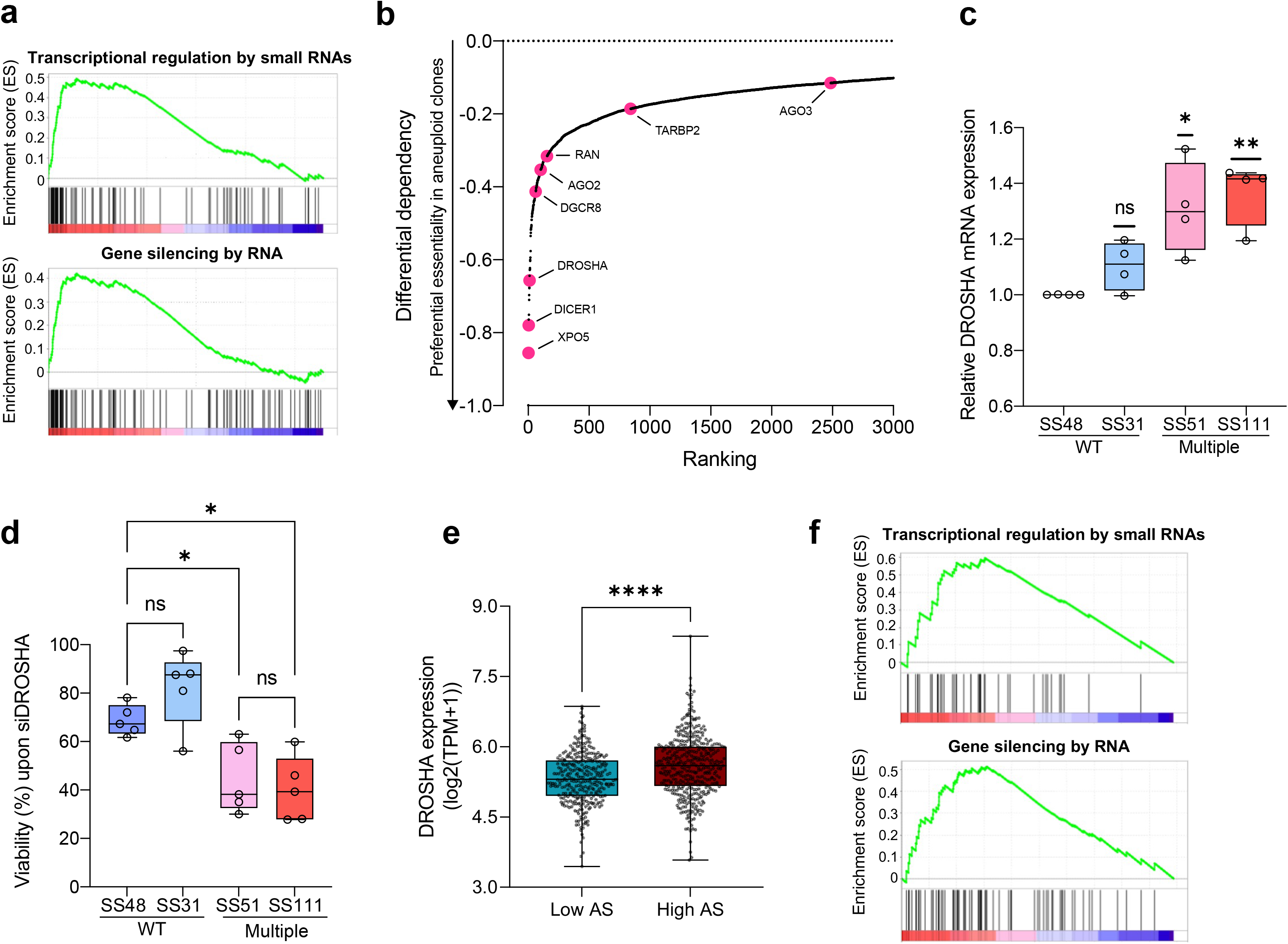
Aneuploid cells activate the miRNA pathway, and depend on this pathway for downregulating their gene expression. **(a)** Gene set enrichment analysis (GSEA) of miRNA-related signatures, comparing the highly-aneuploid clones, SS51 and SS111, to the pseudo-diploid clone SS48. Shown are enrichment plots for the Reactome ‘Transcriptional regulation by small RNAs’ (NES=2.64; q-value<0.0001) and the Reactome ‘Gene silencing by RNA’ (NES=2.36; q-value=0.00016) gene sets. Data are obtained from Zerbib, Ippolito et al, *bioRxiv* 2023. **(b)** The top 3,000 genes that aneuploid clones were most preferentially sensitive to their knockout in comparison to the pseudo-diploid clone SS48, based on our genome-wide CRISPR/Cas9 screen. Highlighted are genes that belong to the miRNA biogenesis pathway (in pink), based on the Reactome ‘miRNA biogenesis’ signature (RNA polymerase II genes excluded). miRNA genes are significantly enriched within the top 3,000 gene list. **, p=0.0064; two-tailed Fisher’s Exact test. Data are obtained from Zerbib, Ippolito et al, *bioRxiv* 2023. **(c)** Comparison of DROSHA mRNA levels, quantified by qRT-PCR, between pseudo-diploid clones (SS48 and SS31) and highly-aneuploid clones (SS51 and SS111). Fold change in DROSHA expression was calculated relative to SS48, per experiment. n=4 independent experiments; *, p=0.0325 and **, p=0.0079, for SS51 and SS111, respectively; One-Sample t-test. **(d)** Comparison of cell viability following siRNA against DROSHA for 72hrs, between pseudo-diploid clones (SS48 and SS31) and highly-aneuploid clones (SS51 and SS111). Viability was calculated relative to control siRNA. n=5 independent experiments; *, p=0.0425 (SS48/SS51) and p=0.0148 (SS48/SS111); One-Way ANOVA, Tukey’s multiple comparison test. All comparisons between SS31 and aneuploid clones were significant as well (**, p<0.01). **(e)** Comparison of DROSHA mRNA expression levels between the top and bottom aneuploidy quartiles of human cancer cell lines (n=738 cell lines). Data were obtained from the DepMap Expression 22Q1 release. DROSHA mRNA expression is significantly higher in highly aneuploid cancer cell lines. ****, p<0.0001; two-tailed Mann-Whitney test. (**f**) Pre-ranked GSEA of mRNA expression levels showing that high aneuploidy levels are associated with upregulation of gene silencing in human primary tumors. Shown are the Reactome ‘Transcriptional regulation by small RNAs’ (NES=1.98; q-value=0.001) and the Reactome ‘Gene silencing by RNA’ (NES=1.86; q-value=0.004) gene sets. Data were obtained from the TCGA mRNA expression data set^62^.

As DROSHA is the most upstream core member of this pathway, we investigated its activity and the sensitivity to its inhibition in the RPE1 clones. The highly-aneuploid clones significantly increased DROSHA mRNA and protein expression (**Fig. 3c, Extended Data Fig. 4a**), and were significantly more sensitive to siRNA-mediated DROSHA depletion, in comparison to the pseudo-diploid clones (**Fig. 3d** and **Extended Data Fig. 4a**). In line with these findings, DROSHA was also significantly over-expressed in highly-aneuploid human cancer cell lines (in comparison to near-euploid human cancer cell lines) (**Fig. 3e**). Whereas aneuploid human cancer cell lines were not more dependent on DROSHA itself, we found that they were more dependent on various other members of the miRNA pathway, and in particular on core members of the RISC complex^34^, such as TRBP (also known as TARBP2) and PACT (also known as PRKRA) (**Extended Data Fig. 4b-c**). Furthermore, high degree of aneuploidy was significantly associated with elevated expression of the miRNA pathway across human primary tumors as well (**Fig. 3f**). Together, these results suggest that miRNA-mediated gene silencing plays an important role in regulating gene expression in aneuploid cells. Interestingly, DROSHA and the miRNA pathway have been previously reported to be involved in DDR^35–38^. We confirmed that etoposide-treated RPE1 cells elevated their DROSHA expression levels (**Extended Data Fig 4d-e**), suggesting a potential role for DDR (Zerbib, Ippolito et al, *bioRxiv* 2023) in mediating the association between aneuploidy and miRNA-mediated RNA degradation.

Lastly, we observed that the aneuploidy-induced changes in RNA metabolism were not limited to RNA degradation – RNA splicing was also among the most differentially-essential pathways in our CRISPR screens (**Fig. 1m**). Thus, we investigated splicing activity in our model system. Several splicing signatures were downregulated in the highly-aneuploid clones (**Extended Data Fig. 4f**), and splicing analysis of the RNAseq data revealed a significant decrease in both 5’ and 3’ alternative splicing in the aneuploid clones (**Extended Data Fig. 4g-h**). These results are consistent with the previously reported competitive interplay between miRNA biogenesis and RNA splicing^39^, further supporting an important role for the miRNA pathway in aneuploid clones.

We conclude that various aspects of RNA metabolism are altered in aneuploid cells, and propose that these cells suffer from transcriptional burden that is offset by increased RNA degradation, making them dependent on the increased activity of two major RNA degradation mechanisms: NMD and miRNAs.

### Increased proteotoxic stress and reduced translation in aneuploid cells

Proteotoxic stress has been reported to be associated with aneuploidy in both yeast^7–10,12^ and engineered aneuploid mammalian cells^2,19,40–43^. Proteotoxic stress may result in reduced protein translation and increased protein degradation, both of which could contribute to dosage compensation at the protein level. Indeed, we identified ongoing proteotoxic stress in our aneuploid clones (**Fig. 1h** and **Extended Data Fig. 1b**). A GSEA analysis comparing the highly-aneuploid clones, SS51 and SS111, and the pseudo-diploid clone SS48, confirmed that highly-aneuploid clones upregulate gene expression signatures of proteotoxic stress and protein degradation (**Fig. 4a**). To validate these results, we characterized the unfolded protein response (UPR) – the primary consequence of proteotoxic stress – in the RPE1 clones. We investigated the three main branches of the UPR^44,45^, and detected increased mRNA expression of two of the canonical UPR branches in highly-aneuploid clones: active XBP1 and EDEM1 indicating elevated activity of the IRE1a branch, and the chaperone BiP (also known as GRP78) indicating elevated activity of the ATF6 branch (**Fig. 4b**), consistent with the RNAseq data (**Fig. 1h** and **Fig. 4a**). We validated the upregulation of GRP78 at the protein level as well (**Fig. 4c-d**). Next, we functionally characterized the UPR in the cells by measuring the response of the isogenic cell lines to the ER stress inducer, tunicamycin^46^. In line with their higher basal level of ER stress, the highly-aneuploid clones, SS51 and SS111, were significantly more resistant to UPR induction (**Fig. 4e**). UPR activation in response to accumulation of misfolded proteins results in translation attenuation^47^. To investigate whether UPR attenuates translation in our model, we performed a SUnSET puromycin incorporation assay^48^. Puromycin incorporation significantly decreased in the highly-aneuploid clones (**Fig. 4f-g**), confirming that global translation levels are reduced in these cells.

**Figure 4:**
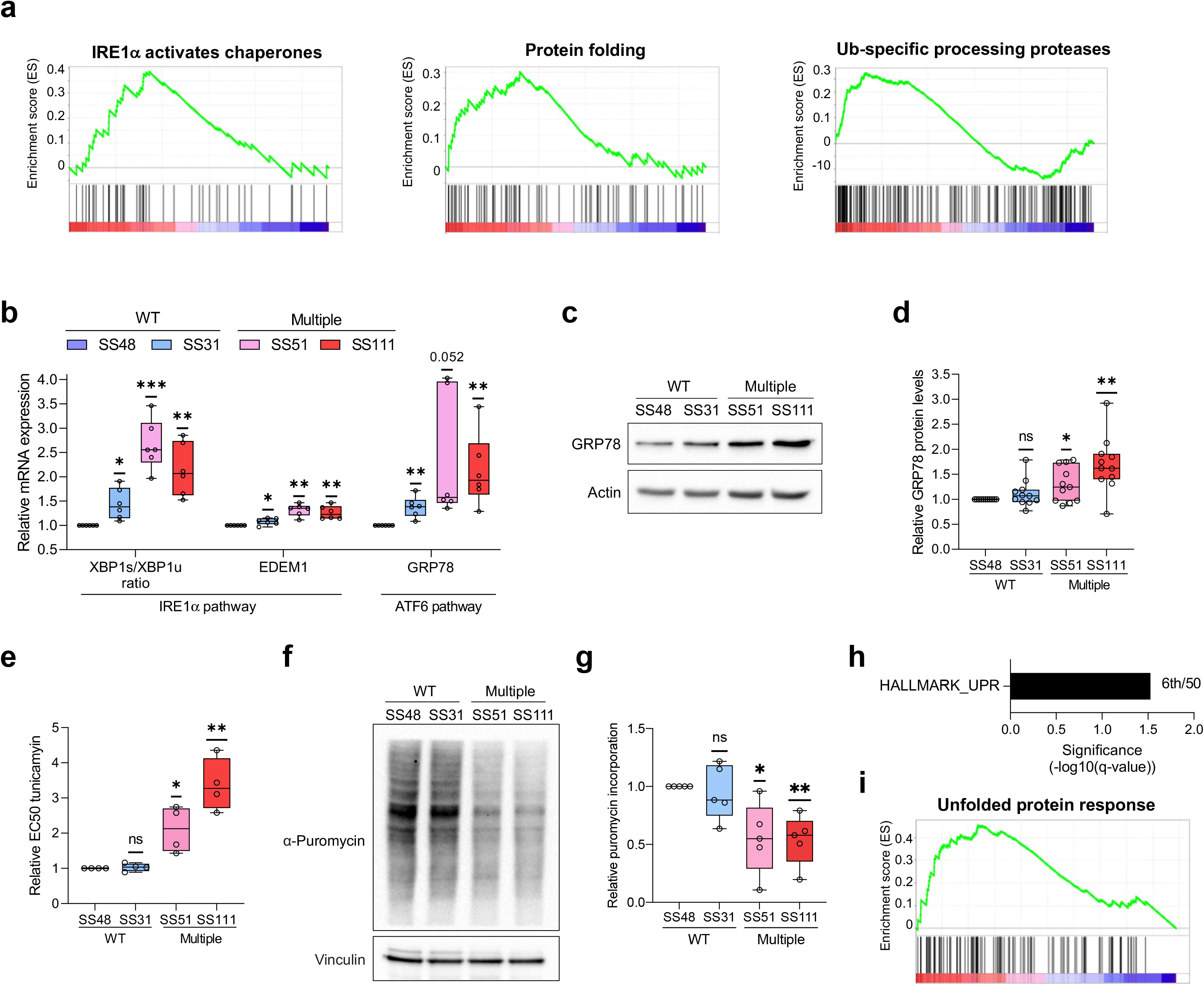
Aneuploid cells experience proteotoxic stress and attenuate protein translation. **(a)** Gene set enrichment analysis (GSEA) of proteotoxic stress-related signatures, comparing the highly-aneuploid clones, SS51 and SS111, to the pseudo-diploid clone SS48. Shown are the enrichment plots for the Reactome gene sets ‘IRE1a activates chaperones’ (NES=1.77; q-value=0.022), ‘Protein folding’ (NES=1.55, q-value=0.084), and ‘Ub-specific processing proteases’ (NES=1.67, q-value=0.041). Data are obtained from Zerbib, Ippolito et al, *bioRxiv* 2023 **(b)** Comparison of UPR mRNA levels, quantified by qRT-PCR, between pseudo-diploid (SS48 and SS31) and highly aneuploid clones (SS51 and SS111). The expression levels of the following canonical members of the UPR were measured: XBP1-spliced/XBP1-unspliced ratio and EDEM1 (IRE1a pathway), and GRP78 (ATF6 pathway). Fold change in expression was calculated relative to SS48, per experiment. n=6 independent experiments; *, p<0.05 and **, p<0.01 in each panel; One-Sample t-test. **(c)** Western blot of GRP78 protein levels in pseudo-diploid clones (SS48 and SS31) and highly-aneuploid clones (SS51 and SS111). β-Actin was used as a housekeeping control. **(d)** Quantification of GRP78 protein levels, calculated relative to SS48 per experiment. n=11 independent experiments; *, p=0.0193 and **, p=0.0019, for SS51 and SS111, respectively; One Sample t-test. **(e)** Comparison of drug sensitivity (determined by EC50 values) to 48hr drug treatment with the UPR activator tunicamycin, between pseudo-diploid clones (SS48 and SS31) and highly-aneuploid clones (SS51 and SS111). EC50 fold-change was calculated relative to SS48, per experiment. n=4 independent experiments; *, p=0.004 and **, p=0.0079, for SS51 and SS111, respectively; One-Sample t-test. **(f)** Representative image of SUNset puromycin incorporation assay, showing reduction in global translation in highly-aneuploid clones (SS51 and SS111) in comparison to pseudo-diploid clones (SS48 and SS31). Vinculin was used as a housekeeping control. **(g)** Quantitative comparison of SUNset puromycin incorporation between pseudo-diploid (SS48 and SS31) and highly-aneuploid clones (SS51 and SS111), calculated relative to SS48. n=5 independent experiments; *, p=0.0323 and **, p=0.009 for SS51 and SS111 respectively; One-Sample t-test. **(h)** Gene set enrichment analysis (GSEA) of the genes whose expression correlates with proliferation in highly-aneuploid cancer cell lines but not in near-diploid cancer cell lines, reveals significant enrichment for UPR. Shown is Hallmark ‘Unfolded Protein Response’. Significance values represent the FDR q-values. The ranking of each proteasome signature (out of all signatures included in the gene set collection) is indicated next to each bar. Data were obtained from DepMap Expression 22Q1 release. **(i)** Pre-ranked GSEA of mRNA expression levels showing that high aneuploidy levels are associated with upregulation of the UPR in human primary tumors. Shown is the Hallmark ‘Unfolded Protein Response’ (NES=1.80, q-value=0.001) gene set. Data were obtained from the TCGA mRNA expression data set^62^.

To assess the generalizability of these findings, we turned to a second isogenic system of RPE1 cells and their aneuploid derivatives, RPTs^49^. In this model, RPE1 cells have doubled their genomes following cytokinesis inhibition, resulting in chromosomal instability and highly-aneuploid cells^49^. The highly-aneuploid RPT cells were indeed more resistant to UPR induction in comparison to their parental pseudo-diploid cells (**Extended Data Fig. 5a**), and exhibited decreased levels of global translation (**Extended Data Fig. 5b-c**). Reversine-mediated aneuploidization of the parental RPE1 cells also resulted in increased resistance to tunicamycin and reduction in global translation (**Extended Data Fig. 5d-f**), further demonstrating that ER stress and reduced translation are an immediate consequence of aneuploidy.

Finally, we investigated the UPR and proteotoxic stress in aneuploid human cancer cells. Gene expression analysis of hundreds of human cancer cell lines showed a significant enrichment for UPR in highly-proliferative highly-aneuploid cancer cell lines (**Fig. 4h**), in line with a recent report^4^. Moreover, a lineage-controlled pan-cancer analysis of The Cancer Genome Atlas (TCGA) mRNA expression datasets revealed a significant elevation of the UPR gene expression signature in highly-aneuploid tumors (**Fig. 4i**), consistent with a recent TCGA analysis that associated UPR with copy number alterations in general^50^. Therefore, we conclude that both non-transformed and cancerous aneuploid cells suffer from proteotoxic stress and must develop compensatory mechanisms to overcome it. One such mechanism is the reduction of the global translation levels, which may be partly responsible for the protein-level dosage compensation observed in aneuploid cells^3,4,19,20,24^.

### Increased proteasome activity and dependency in aneuploid cells

Proteotoxic stress also leads to protein degradation through the ubiquitin-proteasome system^51^. Indeed, our genome-wide gene expression analysis revealed transcriptional signatures of protein degradation to be significantly elevated in the aneuploid clones (**Fig. 1h**), and the proteasome pathway was among the top differential dependencies of aneuploid cells in the CRISPR screen (**Fig. 1m**). We therefore hypothesized that highly-aneuploid cells increase their proteasome activity to overcome proteotoxic stress, and that this makes them more vulnerable to proteasome inhibition. We validated the increased expression and activity of the proteasome complex in the RPE1 models. The highly-aneuploid clones increased the expression of the proteasome subunits (**Fig. 5a**), suggesting an increased proteasome activity in this model. Consistent with this finding, the mRNA expression of the same proteasomal subunits were upregulated in RPT cells (**Extended Data Fig. 6a**), and following reversine treatment of the parental RPE1 cells (**Extended Data Fig. 6b**). Moreover, the highly-aneuploid clones significantly upregulated the chymotrypsin-like activity of their proteasome (**Fig. 5b**), and was similarly observed in RPT cells (**Extended Data Fig. 6c**) and following reversine-induced aneuploidization (**Extended Data Fig. 6d**). Interestingly, the increase in proteasome activity corresponded well with the degree of overexpression of the proteasome subunits across all three model systems. Together, these results suggest that aneuploid cells activate the proteasome system to increase their protein degradation.

**Figure 5:**
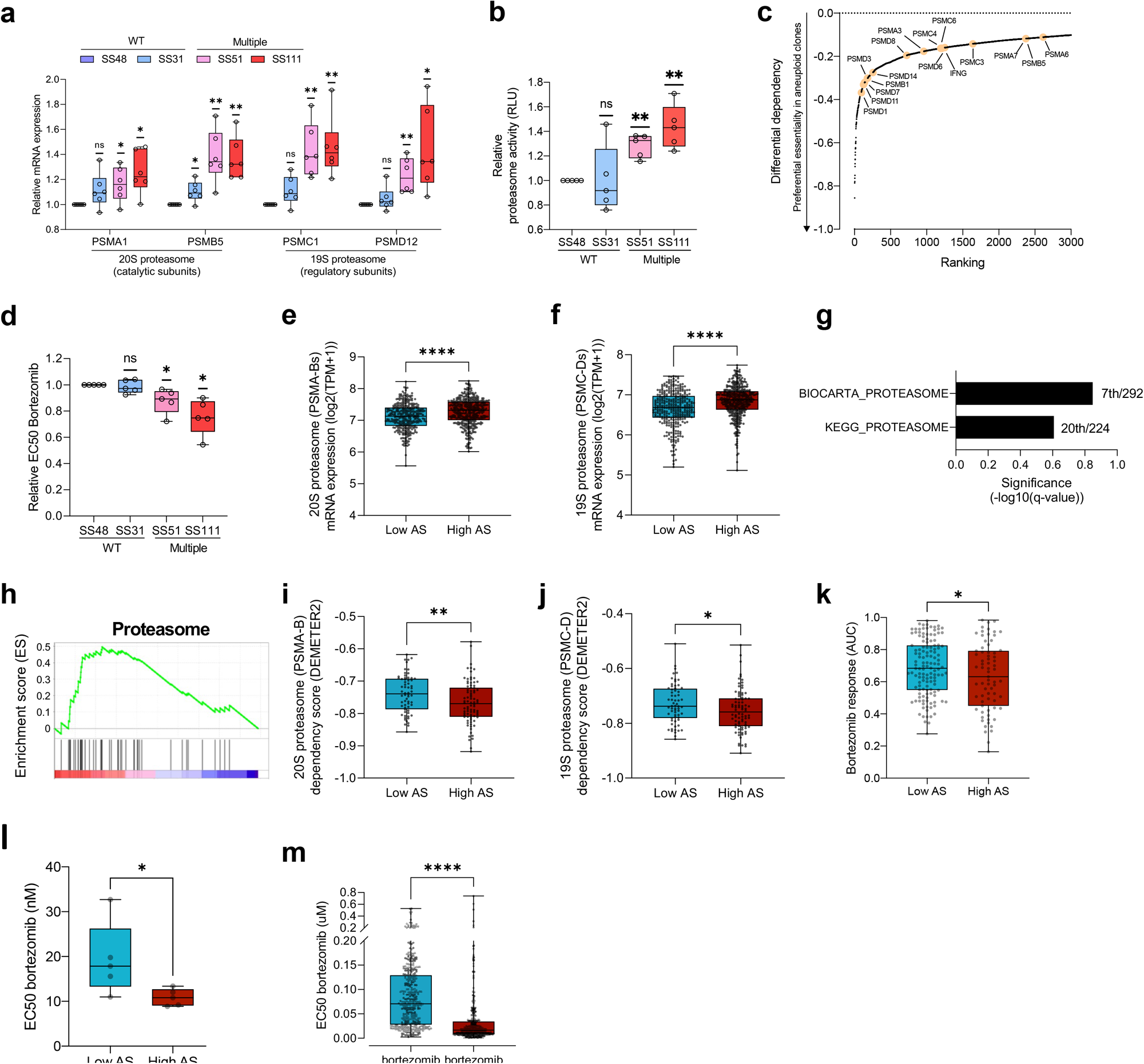
Aneuploid cells activate the proteasome, and depend on its activity for downregulating their protein expression. **(a)** Comparison of mRNA levels, quantified by qRT-PCR, between pseudo-diploid (SS48 and SS31) and highly-aneuploid clones (SS51 and SS111) of representative subunits of the 20S and 19S proteasome complexes: PSMA1, PSMB5, PSMC1, PSMD12. Fold change in expression was calculated relative to SS48, per experiment. n=6 independent experiments; *, p<0.05 and **, p<0.01 in each panel; One-Sample t-test. **(b)** The levels of proteasome activity, measured by Proteasome-Glo®, in pseudo-diploid (SS48 and SS31) and highly-aneuploid clones (SS51 and SS111), showing increased proteasome activity in highly-aneuploid clones. Proteasome activity was calculated relative to SS48, per experiment. n=5 independent experiment, **, p=0.0027 and p=0.0056, for SS51 and SS111 respectively; One-Sample t-test. **(c)** The top 3,000 genes that aneuploid clones were most preferentially sensitive to their knockout in comparison to the pseudo-diploid clone SS48, based on our genome-wide CRISPR/Cas9 screen. Data are obtained from Zerbib, Ippolito et al, *bioRxiv* 2023. Highlighted are genes that belong to the proteasome complex (based on KEGG ‘Proteasome’ gene set). Proteasome genes are significantly enriched within the top 3,000 gene list; *, p=0.0233; two-tailed Fisher’s Exact test. **(d)** Comparison of drug sensitivity (determined by EC50 values) to 72hrs drug treatment with the proteasome inhibitor bortezomib, between pseudo-diploid clones (SS48 and SS31) and highly-aneuploid clones (SS51 and SS111). EC50 fold-change was calculated relative to SS48, per experiment. n=5 independent experiments; *, p=0.0437 and p=0.0163, for SS51 and SS111, respectively; One-Sample t-test. **(e-f)** Comparison of mRNA expression levels of 20S (**e**) and 19S (**f**) proteasome subunits between the top and bottom aneuploidy quartiles of human cancer cell lines (n=738 cell lines). Data were obtained from the DepMap CRISPR screen 22Q1 release. 20S and 19S mRNA expression levels are significantly increased in highly-aneuploid cancer cell lines. ****, p<0.0001; two-tailed Mann-Whitney test. **(g)** Gene set enrichment analysis (GSEA) of the genes whose expression correlates with proliferation in highly-aneuploid cancer cell lines but not in near-diploid cancer cell lines, reveals significant enrichment of proteasome-related signatures. Shown here are Biocarta ‘Proteasome’ and KEGG ‘Proteasome’ signatures. Significance values represent the FDR q-values. The ranking of each proteasome signature (out of all signatures included in the gene set collection) is indicated next to each bar. Data were obtained from DepMap Expression 22Q1 release. **(h)** Pre-ranked GSEA of mRNA expression levels showing that high aneuploidy levels are associated with upregulation of the proteasome in human primary tumors. Shown is the enrichment plot of KEGG ‘Proteasome’ (NES=1.65; q-value=0.042) gene set. Data were obtained from TCGA mRNA expression^62^. **(i-j)** Comparison of gene dependency (determined by DEMETER2 score) for 20S (**i**) and 19S (**j**) proteasome subunits, between the top and bottom aneuploidy quartiles of human cancer cell lines (n=738 cell lines). Data were obtained from the DepMap RNAi screen, 22Q1 release. **, p=0.0089 and *, p=0.0462 for 20S and 19S proteasome subunits, respectively; two-tailed Mann-Whitney test. **(k)** Comparison of drug sensitivity (determined by AUC) to the proteasome inhibitor bortezomib, between the top and bottom aneuploidy quartiles of human cancer cell lines (n=203 cell lines). Data were obtained from GDSC1 drug screen, DepMap portal 22Q1 release. *, p=0.0404; two-tailed t-test test. **(l)** Comparison of drug sensitivity (determined by EC50 values) of 5 near-euploid (CAL51, EN, MHHNB11, SW48 and VMCUB1) and 5 highly-aneuploid (MDA-MB-468, NCIH1693, PANC0813, SH10TC, A101D) cancer cell lines to 72hr drug treatment with the proteasome inhibitor bortezomib. *, p=0.0317; Mann-Whitney test. **(m)** PRISM-based^53^ comparison of drug sensitivity (determined by EC50 values) to 120hr treatment with the proteasome inhibitor bortezomib, between cancer cells treated with the SAC inhibitor reversine (250nM) or with control DMSO (n=387 cell lines). Aneuploidy induction sensitized cancer cells to bortezomib. ****, p<0.0001; two-tailed Wilcoxon rank sum test.

We then turned to investigate the dependency of aneuploid cells on the proteasome. Core proteasomal subunits were among the top differentially-essential genes in the CRISPR screen (**Fig. 5c**), so that aneuploid clones were significantly more sensitive to the perturbation of the 26S proteasome subunits than the pseudo-diploid clone (**Extended Data Fig. 6e**). To validate this finding, we exposed the RPE1 clones to bortezomib, a widely used proteasome inhibitor drug. The highly-aneuploid clones were significantly more sensitive to proteasome inhibition than their pseudo-diploid counterparts (**Fig. 5d**), although the difference was subtle, most likely due to the high potency of the drug. Interestingly, the most aneuploid clone, SS111, also exhibited the strongest UPR response (**Fig. 4e**), the strongest proteasome subunit expression and activity (**Fig. 5a-b**), and the strongest sensitivity to bortezomib (**Fig. 5d**), further supporting the association between aneuploidy and these cellular responses.

Next, we asked whether proteasome activity and dependency are associated with a high degree of aneuploidy in human cancer cells as well. Gene expression analysis of hundreds of human cancer cell lines revealed increased gene expression of both the 20S and 19S proteasome subunits in highly-aneuploid cancer cells (**Fig. 5e-f**). Moreover, genes associated with the proliferation capacity of highly-aneuploid, but not of near-euploid, cancer cell lines were strongly enriched for proteasome signatures (**Fig. 5g**). Importantly, we found a significant association between aneuploidy and the proteasome gene expression signature in the TCGA dataset as well (**Fig. 5h**), suggesting that this association holds true in primary tumors. Together, these results suggest an increased proteasome activity in highly-aneuploid cancer cells.

Finally, we investigated the association between aneuploidy and proteasome dependency in human cancer cells. Highly-aneuploid cancer cells were more dependent on genetic (shRNA-mediated) silencing of both the 20S and 19S proteasome subunits (**Fig. 5i-j**) and more sensitive to pharmacological inhibition using bortezomib (**Fig. 5k**). To validate these findings, we selected five representative cancer cell lines with low degree of aneuploidy and five representative cancer cell lines with a high degree of aneuploidy^52^, and compared their response to bortezomib. Indeed, highly-aneuploid cancer cells were more sensitive to the proteasome inhibitor (**Fig. 5l** and **Extended Data Fig. 6f-g**). To confirm that proteasome dependency is indeed causally related to aneuploidy in cancer cells, we assessed the response of 578 human cancer cell lines to bortezomib, using the PRISM barcoded cell line platform^53^. The response to bortezomib was evaluated either in the absence or in the presence of a low dose (250nM) of reversine (see **Methods**). At this concentration, reversine had a mild effect on proliferation (**Supp. Table 1**), but significantly sensitized cancer cells to proteasome inhibition (**Fig. 5m**). Therefore, we conclude that aneuploid cancer cells upregulate their proteasome activity in response to proteotoxic stress, rendering them more sensitive to proteasome inhibition.

## Discussion

### RNA metabolism in aneuploid cells

In all organisms analyzed to date, changes in gene copy number generally trigger corresponding changes in the amount of produced mRNA^7–9,12,17,19,54–56^. In agreement with this, our data show that aneuploid cells experience increased RNA synthesis (**Fig. 1**). Importantly, we also found that aneuploid cells upregulate pathways involved in RNA degradation and gene silencing, such as the nonsense-mediated decay and the miRNA pathways (**Fig. 2** and **Fig. 3**). These data argue that buffering mechanisms might be at play in aneuploid cells to limit the burden brought about by an imbalanced karyotype. Although this stoichiometric control has been extensively studied at the protein level in the context of aneuploidy — in both untransformed^7–10,12,17,19,55–57^ and cancer cells^4,5^ – the role and the impact of RNA metabolism regulation in controlling gene expression is only beginning to emerge as another important layer of regulation^23,58,59^. Interestingly, dosage compensation at the mRNA level seems to be minimal in yeast^13,20^, but has been recently observed in human cells^4,6^.

Our findings of elevated gene silencing pathways in aneuploid cells indicate the existence of a dynamic regulation of gene expression, which, most likely, acts in concert with the regulation of protein homeostasis of multimeric complexes, and could also be facilitated by post-translational modifications^3,9^. Intriguingly, the effect of aneuploidy on RNA metabolism is not limited to the gained chromosomes or to protein complex genes, as we did not observe increased RNA degradation of transcripts from such genes. These findings are consistent with recent reports that dosage compensation of protein complex genes mostly occurs at the protein regulation level^5,6^. How aneuploid cells evolve to alter their global RNA metabolism in response to changes in gene dosage remains to be fully understood. In this respect, there are at least two possible scenarios: gene silencing activities might be the direct consequence of increased gene expression, somehow sensed by the cells; or could be induced indirectly following aneuploidy-induced cellular stresses. We favor the latter possibility and speculate that a major aneuploidy-induced stress playing a role in this process is DNA damage. Indeed, our data show an increased expression of the NMD core component CASC3 following DNA damage induction in pseudo-diploid RPE1 cells, in full agreement with previous reports indicating that DDR triggers NMD activity^29–32^. We propose that aneuploidy-induced cellular stresses result in altered RNA metabolism in aneuploid cells, counteracting changes in gene expression caused by imbalanced karyotypes.

Importantly, the increased dependency of the aneuploid cells on RNA degradation mechanisms was independent of p53 status (**Extended Data Fig. 1**), indicating that this is a consequence of the aneuploid state *per se*. We note, however, that our isogenic cell lines harbored extra chromosomes (trisomies), and the dosage compensation mechanisms that we identify are therefore associated with trisomies rather than with aneuploidy in general; different mechanisms for dosage compensation may be triggered upon monosomy^4,60^, and should be specifically addressed in future studies.

### Proteotoxic stress and proteasome dependency in aneuploid cells

Tight control of pathways involved in protein translation and degradation is crucial to limit proteotoxic stress in aneuploid cells^2,7–10,12,17,19,55,57^. Proteotoxic stress is perhaps the most prominent consequence of karyotype imbalances; the simultaneous overexpression of hundreds or thousands of genes, brought about by gained chromosomes, lead to a massive burden on protein homeostasis. The effects of aneuploidy-induced proteotoxic stress that have been described so far are mainly: (1) the overwhelming of the protein-folding machineries^2,40,61^ ; and (2) the saturation of catabolic pathways responsible for the degradation of excessive proteins^9,17,19,40^. Importantly, our results indicate that aneuploid cells are sensing and responding to the altered demand in the synthesis, folding and assembly of proteins both by attenuating global protein translation and by reducing global protein degradation (**Fig. 4**), thereby “buffering” the stoichiometric imbalance induced by aneuploidy.

Interestingly, recent studies have shown that protein buffering is common in cancer cells, and suggest that maintenance of proper protein complex stoichiometries is crucial for tumor growth^30,31^. A recent analysis of TCGA data revealed that the abundance of the proteasome subunits was correlated with the degree of stoichiometric imbalance. In the current study, we took this notion further, demonstrating that aneuploid cancer cells not only activate the proteasome but also become more dependent on its activity, making them more sensitive to proteasome inhibition (**Fig. 5**). We propose that aneuploidy might be a biomarker for predicting tumor’s response to proteasome inhibitors.

## Concluding remarks

Extensive transcriptome and proteome imbalance is one of the most immediate and important consequences of aneuploidy. Our work indicates that RNA and protein metabolism – and in particular their degradation – play a central role in attenuating the cellular impact of the increased DNA content that inevitably characterizes trisomic cells. Therefore, dosage compensation might be achieved by perturbation of various stages along the gene expression process (**Fig. 6**). Importantly, each of these stages presents a potential opportunity for therapeutic intervention: cardiac glycosides might represent a novel class of anti-aneuploid cancer therapeutics through targeting of NMD; and proteasome inhibitors might be preferentially effective against aneuploid cancer cells due to their increased reliance on the proteasome activity (**Fig. 6**). As these drugs are already used in the clinic, clinical trials are now necessary to determine if they can indeed be used to treat aneuploid tumors.

**Figure 6:**
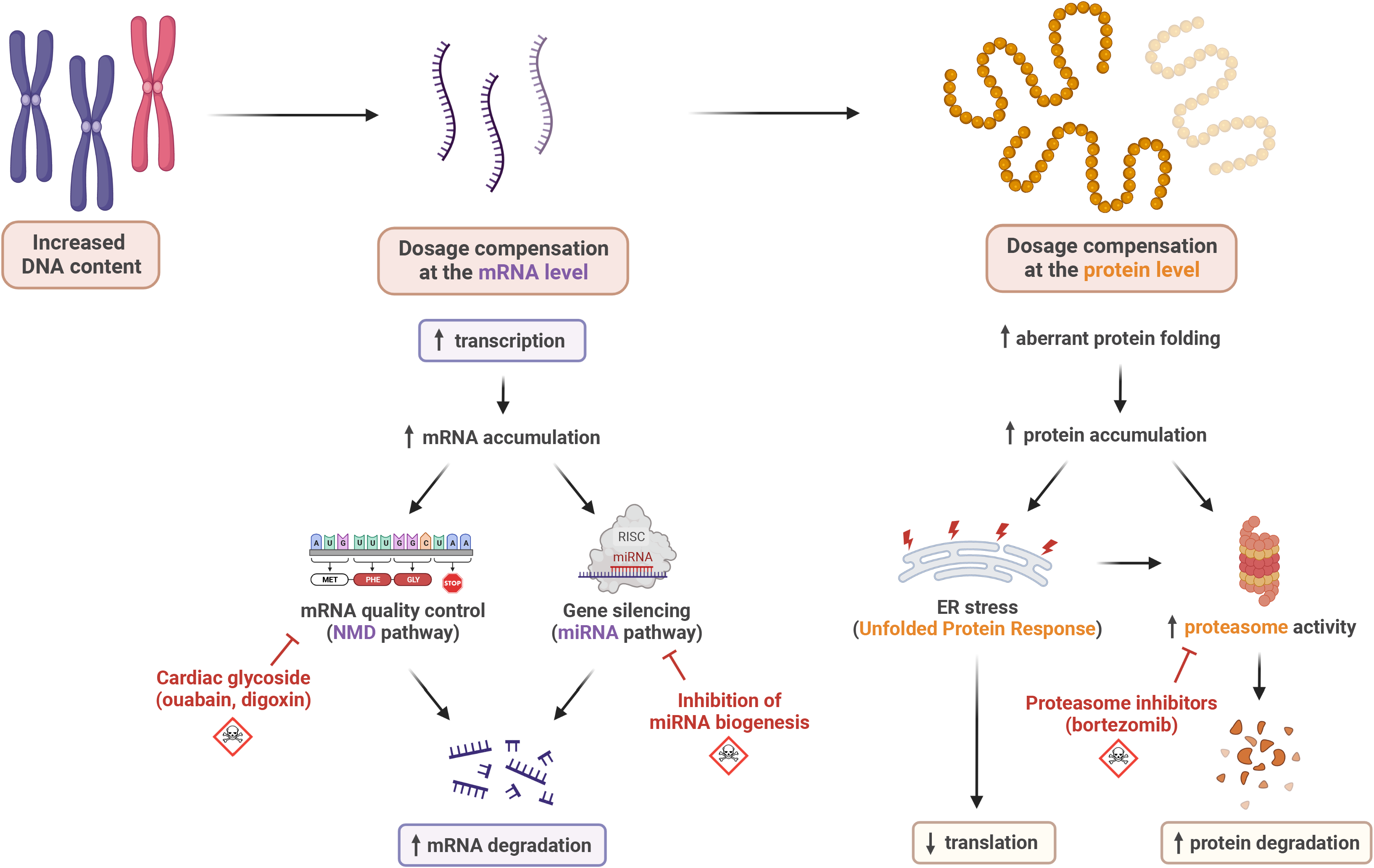
Aneuploid cells with extra chromosomes compensate for their excessive DNA content at both the RNA and the protein level. A summary illustration of the study. Increased DNA content leads to increased transcription in aneuploid cells, which is counterbalanced by reducing the cellular mRNA levels via activation of the NMD and the miRNA pathways. The increase in the number of total and aberrant transcripts induces accumulation of misfolded proteins that triggers the UPR. Consequently, aneuploid cells decrease their protein translation and increase their protein degradation by activating the proteasome machinery. Aneuploid cells therefore become preferentially sensitive to the perturbation of both RNA and protein metabolism.

## Methods

### Cell culture

RPE1-hTERT cells, their derivatives clones and RPT, CAL51, EN, VMCUB, SW48, MDA-MB-468 and A101D cell lines, were cultured in DMEM (Life Technologies) with 10% fetal bovine serum (Sigma-Aldrich), 1% sodium pyruvate, 4mM glutamine, and 1% penicillin-streptomycin. SH10TC, NCIH1693, MHHNB11 and PANC0813 were cultured in RPMI-1640 (Life Technologies) with 10% fetal bovine serum (Sigma-aldrich) and 1% penicillin-streptomycin-glutamine (Life Technologies). PANC0813 medium was supplemented with 10units/mL human recombinant insulin (Sigma-Aldrich), and MHHNB11 medium was supplemented with MEM Non-Essential Amino Acids (Sigma-Aldrich). Cells were cultured at 37°C with 5% CO2 and are maintained in culture for maximum three weeks. All cell lines were tested free of mycoplasma contamination using Myco Alert (Lonza, Walkersville, MD, USA) according to the manufacturer’s protocol. To induce random aneuploidy, cells were seeded and synchronized with 5mM Thymidine for 24hrs, then treated with 500nM reversine (or vehicle control) for 16hrs. Read-outs were performed 72hrs post reversine wash-out.

### RNA synthesis

Cells were seeded on coverslips coated with 5μg/ml fibronectin. 72hrs later, EZClick™ RNA label was incubated for 1h at 37°C. Then, De novo synthesized RNA and DAPI were detected following manufacturer’s instructions. Coverslips were mounted using Mowiol. Cells were imaged using Leica SP8 confocal microscope with a magnification objective of 40x. FIJI software was used for the quantification of nascent RNA spots area.

### RNAseq and data analysis

RNA sequence reads were obtained from and were analyzed as previously described (Zerbib, Ippolito et al, *bioRxiv* 2023). Normalized read counts, and differential gene expression analysis were generated using DESeq2 R package^63^. GSEA and pre-ranked GSEA were performed on the differentially expressed genes using GSEA software 4.0.3, with the following parameters: 1000 permutations and Collapse analysis, using the Hallmark, KEGG, Biocarta, and Reactome gene sets (in separate analyses). Genes with fewer than 10 and 20 normalized read counts, for GSEA and pre-ranked GSEA respectively, were excluded from further analyses. Evaluation of degraded RNA was performed using Degnorm with default parameters, as previously described^25^, to generate the degradation index (DI) and the degradation-free expression matrix. GSEA was then repeated with the degradation-free expression matrix.

NMD pathway transcriptional activity was evaluated as previously described^26^. Briefly, we calculated the R_mRNA_ score, i.e. the mRNA abundance of an NMD target gene, following the equation: R_mRNA_ = mE_NMD_ /median_mE_non-NMD_ (mE_NMD_ being the mRNA expression of the NMD target, and median_mE_non-NMD_ being the median of mRNA expression of non-NMD target genes). To infer the NMD pathway activity in aneuploid clones, an NMD transcriptional score, representing the relative abundance of the NMD target gene in aneuploid clones compared to pseudo-diploid RPE1-SS48, was calculated following the equation: NMD score=R_mRNA_ (aneuploid)/R_mRNA_ (SS48).

Differential splicing analysis was performed using VAST-Tool^64^. RNAseq reads were aligned against the VASTDB of the human reference genome hg19. The Percent Spliced-In (PSI) score for each splicing event, representing the percentage of included splicing events out of total splicing events (higher the index, lower the splicing activity), was calculated using the Vast-tool package and “compare” method, between SS48 and each one of the aneuploid samples. For the downstream analysis, only the alternative 3’/5’ splice site events (Alt3, Alt5) with PSI>5 were considered.

### Total RNA electrophoresis

RNA was harvested from 1 million cells using Bio-TRI® (BioLabs) following the manufacturer’s protocol. RNA was run in 1% agarose gel in a cleaned chamber, and migration was imaged every 20min. Smear quantification was performed using ImageJ, by quantifying the smear between the 28S and 16S bands, relative to the total amount of RNA.

### Genome-wide CRISPR screens and data analysis

CRISPR dependency scores (CERES scores) were obtained from Zerbib, Ippolito et al, *bioRxiv* 2023 Dependency analysis was performed as previously described (Zerbib, Ippolito et al, *bioRxiv* 2023), by a pre-ranked GSEA was on the differentially-expressed genes using GSEA software 4.0.3, with the following parameters: 1,000 permutations and Collapse analysis, using the Hallmark, KEGG, Biocarta, and Reactome gene sets (in separate analyses).

### Dependency Map data analysis

Extension of the aneuploidy scores (AS) table of each cancer cell line was obtained from (Zerbib, Ippolito et al, *bioRxiv* 2023). mRNA gene expression values, CRISPR and RNAi dependency scores (Chronos and DEMETER2 scores, respectively) were obtained from DepMap 22Q1 release (https://figshare.com/articles/dataset/DepMap_22Q1_Public/19139906), and compared between the bottom (AS≤ 8) and top (AS≥21) aneuploidy quartiles.

Doubling time (DT) analyses was performed as previously described (Zerbib, Ippolito et al, *bioRxiv* 2023). Briefly, using the extended aneuploidy score table, and within the bottom (AS≤ 8) and the top quartile (AS≥21), DT of each cancer cell line^65^ was correlated to gene expression utilizing a linear model following the method of Taylor *et al*^66^. Genes were determined as overexpressed in highly proliferative aneuploid cancer cells if they were significantly associated with DT within the top AS quartile but not within the bottom AS quartile. Significance thresholds: (log10(p-value)≥2.5) OR (–log10(p-value)≥1.3 AND correlation coefficient<-0.005). The resultant list of genes is available as supplementary table in (Zerbib, Ippolito et al, *bioRxiv* 2023). This list was subjected to gene set enrichment analysis using the ‘Hallmark’, ‘KEGG’, ‘Reactome’ and ‘Gene Ontology Biological Processes’ gene set collections from MSigDB (http://www.gsea-msigdb.org/gsea/msigdb/)^22,67^.

### qRT-PCR

Cells were harvested using Bio-TRI® (Bio-Lab) and RNA was extracted following manufacturer’s protocol. cDNA was amplified using GoScript™ Reverse Transcription System (Promega) following manufacturer’s protocol. qRT-PCR was performed using Sybr® green, and quantification was performed using the ΔCT method. All primer sequences are available in **Supplementary Table 2**.

### NMD pathway reporter assay

NMD pathway reporter assay was performed as previously described^68^. Briefly, 300,000 cells were seeded in 6-well plates and transfected 24hrs later with 2ug of pBS-(CBR-TCR(PTC))-(CBG-TCR(WT)) plasmid^68^ using *Trans*IT-LT1® (Mirus, MIR2300), following manufacturer’s protocol. Medium was replaced 24hrs post-transfection. 72hrs post-transfection, RNA was harvested from the treated cells using Bio-TRI® (BioLabs) following the manufacturer’s protocol. RNA was cleaned from plasmid contamination using TURBO DNA-free™ Kit (Invitrogen, AM1907) following the manufacturer’s protocol. cDNA was amplified using GoScript™ Reverse Transcription System (Promega) following the manufacturer’s protocol. qRT-PCR was performed using Sybr® green, and quantification was performed as previously described^68^.

### Drug treatments

Drug treatments were performed as previously described (Zerbib, Ippolito et al, *bioRxiv* 2023). Briefly, cells were seeded in a 96w plate using Multidrop™ Combi Reagent Dispenser (ThermoFisher), then treated 24hrs later with drugs of interest. Cell viability was measured at indicated time point using the MTT assay (Sigma M2128). Formazan crystals were extracted using 10% Triton X-100 and 0.1N HCl in isopropanol, and color absorption was quantified at 570nm and 630nm. EC50 for each drug was calculated using GraphPad PRISM 9.1, inhibitor vs. response (four parameters) non-linear regression model.

Validation of bortezomib treatment was performed on 5 near-euploid (CAL51, EN, MHHNB11, SW48 and VMCUB1) and 5 highly aneuploid (MDA-MB-468, NCIH1693, PANC0813, SH10TC, A101D) cancer cell lines. Cells were seeded in a 96w plate, and treated 24hrs later with various concentrations of bortezomib. Cell viability was measured after 72hrs using CellTiter-Glo (Promega). EC50 was calculated using GraphPad PRISM 8, asymmetric (five parameters) non-linear regression model. In **Extended Data Fig. 6g**, Cal51 and MDA-MB-468 were imaged after 72hrs exposure to bortezomib, using Incucyte (Satorius). For visualization, the cell borders were highlighted using AI-trained Ilastik® software. All drug details are available in **Supplementary Table 2**.

### siRNA transfection

Cells were transfected with siRNAs against CASC3 or DROSHA (ONTARGETplus SMART-POOL®, Dharmacon), or with a control siRNA (ONTARGETplus SMART-POOL®, Dharmacon) using Lipofectamine® RNAiMAX (Invitrogen) following manufacturers’ protocols. To test whether aneuploidy induction sensitized cells to CASC3, cells were seeded and synchronized with Thymidine 5mM for 24hrs, then treated with reversine 500nM for 20hrs. After the reversine pulse, cells were reverse transfected with siRNA against CASC3 using Lipofectamine® RNAiMAX following the manufacturer’s protocol. Cell growth following siRNA transfection was followed by live cell imaging using Incucyte® (Satorius). The effect of the knockdown on viability was calculated by comparing the cell number in the targeted siRNA vs. control siRNA wells at 72hrs post transfection.

### Western blot

Cells were lysed in NP-40 lysis buffer (1% NP-40;150mM NaCl; 50mM Tris HCl pH 8.0) with the addition of protease inhibitor cocktail (Sigma-Aldrich #P8340) and phosphatase inhibitor cocktail (Sigma Aldrich #P0044). Protein lysates were sonicated (Biorector) for 5min (30sec on/30sec off) at 4° c, then centrifuged at maximum speed for 15 min and resolved on 12% SDS-PAGE gels. Bands were detected using chemoluminescence (Millipore #WBLUR0500) on Fusion FX gel-doc (Vilber). For SUnSET puromycin incorporation assay, cells were treated with 10µg/mL puromycin for 30min prior to harvest. All antibodies are listed and their use is described in **Supplementary Table 2**.

### Proteasome activity assay

Proteasome activity was estimated using Proteasome Glo® Chemotrypsin-like kit (Promega) following manufacturer’s protocol. Briefly, cells were trypsinized and washed twice with medium to remove residual trypsin. 4,000 cells were seeded in triplicate in a white 96-well plate, and incubated for 2hrs at 37°C. 30min exposure to 1µM of bortezomib was used as a positive control for proteasome activity inhibition. Plate was shaken for 2min at high speed, incubated for 5min at RT, and luminescence was then measured using a Synergy H1 plate reader (BioTEK).

### PRISM screen

PRISM screen was performed as previously described^52,53^. Briefly, cells were plated in triplicate in 384-well plates at 1,250 cells per well. Cells were treated with the proteasome inhibitor bortezomib (8 concentrations of threefold dilutions, ranging from 91nM to 20µM) in presence of reversine (250nM) or DMSO for 5 days. Cells were then lysed, and lysate plates were pooled for amplification and barcode measurement. Viability values were calculated by taking the median fluorescence intensity of beads corresponding to each cell line barcode, and normalizing them by the median of DMSO control. Dose-response curves and EC50 values were calculated by fitting four-parameter curves to viability data for each cell line, using the R drc package^69^, fixing the upper asymptote of the logistic curves to 1. EC50 comparisons were performed on the 387 cell lines for which well-fit curves (r^2^ >0.3) were generated.

### TCGA data analysis

TCGA data were retrieved using TCGAbiolinks R package^62^. Aneuploidy scores (AS) were obtained from Taylor *et al*^66^, and correlated to tumor gene expression using lineage as a covariate (lm function in R studio v4.1.1, using the equation: gene∼AS+lineage), as previously described^66^. Genes were ranked based on their aneuploidy score coefficient, and then subjected to pre-ranked gene set enrichment analysis^22^ using the ‘Hallmark’, ‘Biocarta’, ‘KEGG’, and ‘Reactome’ gene set collections from MSigDB.

### Statistical analyses

The number of cells used for each experiment is available in the method section. Western Blot quantifications were performed using ImageJ® and Image Lab. The numbers of independent experiments and analyzed cell lines of each computational analysis are available in the figure legends. Statistical analyses were performed using GraphPad PRISM® 9.1. Details of each statistical test are indicated in the figure legends. In each presented box plot, the internal bar represents the median of the distribution. In **Fig. 1c** and **Fig. 1f**, the bar represents the mean and SEM. Significance thresholds were defined as p-value = 0.05 and q-value = 0.25.

## Acknowledgments

The authors would like to thank James McFarland and Ofir Hamieri for their bioinformatic support; Gil Ast, Marina Mapelli, Zuzana Tothova and members of the Ben-David and Santaguida labs for helpful discussions; Varda Wexler for assistance with Figure preparation; Zuzana Storchova for providing the RPE1/RPT cell lines; Zhongsheng You for providing the NMD reporter constructs; and Nicholas Lyons, Jordan Bryan, Samantha Bender and Jennifer Roth for their assistance with the PRISM screen. We thank the Broad Institute Genomic Perturbation Platform for their assistance with the CRISPR/Cas9 screens. This work was supported by the European Research Council Starting Grant (grant #945674 to U.B.-D.), the Israel Cancer Research Fund Gesher Award (U.B.-D.), the Azrieli Foundation Faculty Fellowship (U.B.-D.), the DoD CDMRP Career Development Award (grant #CA191148 to U.B.-D.), the Israel Science Foundation (grant #1339/18 to U.B.-D.), the BSF project grant (grant #2019228 to U.B.-D.), the Italian Association for Cancer Research (AIRC-MFAG 2018 - ID. 21665 project to S.S.), Ricerca Finalizzata (GR-2018-12367077 to S.S.), Fondazione Cariplo (S.S.), the Rita-Levi Montalcini program from MIUR (to S.S.) and the Italian Ministry of Health with Ricerca Corrente and 5×1000 funds (S.S.). U.B.-D. is an EMBO Young Investigator. J.Z. was supported by a fellowship of the Israeli Ministry for Immigrant Absorption and by travel awards from the TAU Constantiner Institute and Cancer Biology Research Center. M.R.I. is supported by an AIRC Fellowship (ID 26738-2021). J.Z, Y.E, and G.L. are PhD and MD-PhD students within the graduate school of the Faculty of Medicine, Tel Aviv University. M.R.I., S.M, S.V. and S.S. are PhD students within the European School of Molecular Medicine (SEMM).

## Extended Data Figure Legends

**Extended Data Figure 1: Increased RNA degradation and dependency to RNA metabolism is independent to p53-mutation status (related to Fig. 1) (a)**: The fractions of significantly upregulated and downregulated genes out of all differentially-expressed genes, between the highly-aneuploid clones, SS51 and SS111, and the pseudo-diploid clones SS48 and SS77 (qvalue<0.25). n=3006 genes. ****, p<0.0001; two-tailed binomial test. Data are taken from Zerbib, Ippolito et al, *bioRxiv* 2023. **(b)** Comparison of the differential gene expression patterns (pre-ranked GSEA results) between the near-diploid SS48 clone (control) and the aneuploid SS6, SS119, SS51 and S111 clones. Plot presents enrichments for the Hallmark, KEGG, Biocarta and Reactome gene sets. Significance threshold set at qvalue=0.25. Enriched pathways are color-coded. Data are taken from Zerbib, Ippolito et al, *bioRxiv* 2023 **(c)** Comparison of the differential gene dependency scores (pre-ranked GSEA results) between the near-diploid SS48 and SS77 clones (control) and the aneuploid SS6, SS119 and SS51 clones. Data are taken from Zerbib, Ippolito et al, *bioRxiv* 2023. Plot presents enrichments for the Hallmark, KEGG, Biocarta and Reactome gene sets. Significance threshold set at qvalue=0.25. Enriched pathways are color-coded.

**Extended Data Figure 2: RNA degradation does not confound the results of the differential gene expression analyses (related to Fig. 1) (a)**Comparison of the differential gene expression patterns (pre-ranked GSEA results) between the near-diploid SS48 clone (control) and the highly-aneuploid SS51 and SS111 clones, following Degnorm normalization. The plot presents enrichments for the Hallmark, KEGG, Biocarta, and Reactome gene sets. Significance threshold set at qvalue=0.25. Enriched pathways are color-coded. **(b-d)** Gene Set Enrichment Analysis (GSEA) between the near-diploid SS48 clone (control) and the highly-aneuploid SS51 and SS111 clones following Degnorm normalization. The top row **(b)** presents signatures related to DNA damage response and p53 pathway, middle rows **(c)** presents signatures related to RNA metabolism, and bottom row **(d)** presents signatures related to protein metabolism. Presented signatures:

- Reactome ‘Non-homologous End Joining’: NES=2.37; q-value=0.0002
- Reactome ‘Homology Direct Repair’: NES=1.58; q-value=0.034
- Reactome ‘DNA double strand break response’: NES=2.51; q-value=0.0004
- Reactome ‘Base Excision Repair’: NES=3.15; q-value<0.0001
- CPG ‘Kannan TP53 Targets Up’: NES=2.16; q-value=0.0049
- GO-Biological Processes ‘Negative regulation of RNA catabolic process’: NES= -1.42; q-value=0.295
- Reactome ‘Nonsense Mediated Decay’: NES=1.21; q-value=0.19
- GO-Biological Processes ‘Nuclear-transcribed mRNA catabolic processes NMD’: NES=1.32; q-value=0.28
- Reactome ‘Transcriptional regulation by small RNAs’: NES=2.82; q-value<0.0001
- Reactome ‘Gene silencing by RNA’: NES=2.55; q-value=0.0004
- GO-Biological Processes ‘Regulation of RNA splicing’: NES=-1.74; q-value=0.194
- GO-Biological Processes ‘Regulation of mRNA splicing via spliceosome’: NES=-1.75; q-value=0.184
- GO-Biological Processes ‘Regulation of alternative splicing via spliceosome’: NES=-1.72; q-value=0.1998
- Reactome ‘E3-Ub ligase ubiquitinate target proteins’: NES=1.529; q-value=0.042
- Reactome ‘Protein ubiquitination’: NES=1.43; q-value=0.07

**Extended Data Figure 3: Aneuploid cells activate the nonsense-mediated decay (NMD) pathway, and depend on it for downregulating their gene expression (related to Fig. 2) (a)** Gene set enrichment analysis (GSEA) of the nonsense-mediated decay (NMD) gene expression signature, comparing the highly-aneuploid clones SS51 and SS111, to the pseudo-diploid clone SS48. Shown is the enrichment plot for Reactome ‘Nonsense-mediated decay’ gene set (NES=1.91; q-value=0.0085). Data are obtained from Zerbib, Ippolito et al, *bioRxiv* 2023 **(b)** Comparison of the NMD pathway activity between highly-aneuploid clones (SS51 and SS111) and pseudo-diploid clones (SS48 and SS31), using an NMD reporter assay^27,68^. The abundance of the triggering CBR-TCR sequence (PTC) relative to that of the normal CBG-TCR sequence (WT) was measured by qRT-PCR 72hrs post-transfection. Fold change in the PTC/WT ratio was calculated relative to SS48, per experiment. Decreased PTC/WT ratio reflects the high degradation rate of the triggering sequence over the internal control, showing increased NMD pathway activity. n=7 independent experiments; **, p=0.0046 and ***, p=0.0002, for SS111 and SS51, respectively; One-Sample t-test. **(c)** The top 3,000 genes that aneuploid clones were most preferentially sensitive to their knockout in comparison to the pseudo-diploid clones SS48, based on our genome-wide CRISPR/Cas9 screen. Highlighted are genes that belong to the NMD pathway: core member genes (in pink) and ribosomal-related genes (in purple). NMD genes are significantly enriched within the top 3,000 gene list. ****, p<0.0001; two-tailed Fisher’s Exact test. Data are taken from Zerbib, Ippolito et al, *bioRxiv* 2023. **(d)** Comparison of drug sensitivity (determined by IC50 values) to 72hr drug treatment with the NMD inhibitor digoxin, between pseudo-diploid clones (SS48 and SS31) and highly-aneuploid clones (SS51 and SS111). IC50 fold-change was calculated relative to SS48, per experiment. n=4 independent experiments; *, p=0.0207 and **, p=0.0021, for SS111 and SS51, respectively; One-Sample t-test. **(e)** Western blot of CASC3 protein levels in inducible *TP53*-KD RPE1 parental cells, pre-treated with the SAC inhibitor reversine (500nM) or with control DMSO for 20hrs to induce aneuploidy, then harvested 72hrs post wash-out. Vinculin was used as housekeeping control. **(f)** Quantification of CASC3 protein levels in the reversine-treated *TP53*-KD RPE1 parental cells, calculated relative to the DMSO control per experiment. n=3 independent experiments. **, p=0.0096 and *, p=0.0167 for reversine-treated sh-CTL and sh-p53 respectively; One Sample t-test. **(g)** Western blot of CASC3 protein levels in *TP53*-KO RPE1 parental cells, pre-treated with the SAC inhibitor reversine (500nM) or with control DMSO for 20hrs to induce aneuploidy, then harvested 72hrs post wash-out. Vinculin was used as housekeeping control. **(h)** Quantification of CASC3 protein levels in the reversine-treated *TP53*-KO RPE1 parental cells, calculated relative to the DMSO control per experiment. n=5 independent experiments. *, p=0.0185 and ***, p=0.0005 for reversine-treated sg-CTL and sg-p53 respectively; One Sample t-test. **(i)** Western blot of CASC3 protein levels in RPE1 clones treated with siRNA against CASC3 (or control siRNA) for 72hrs. Tubulin was used as a housekeeping control. **(j)** Western blot of CASC3 protein levels in reversine-treated parental RPE1 cells, treated with siRNA against CASC3 (or control siRNA) for 72hrs. Tubulin was used as a housekeeping control. **(k)** Western blot of CASC3 and γH2AX protein levels in parental RPE1 cells treated with etoposide (2.5µM) for 1, 3 or 6 hours. Elevation of CASC3 protein levels is associated with the degree of DNA damage in the cells. Tubulin was used as a housekeeping control. **(l)** Quantification of CASC3 protein levels in parental RPE1 etoposide-treated cells, calculated relative to the DMSO control per experiment. n=5 independent experiments. *, p=0.0311, p=0.0184, p=0.0171 for 1h, 3h, and 6h etoposide treatment respectively; One Sample t-test. **(m-p)** Comparison of gene dependency (determined by Chronos score) for key members of the NMD pathway, EJC member EIF4A3 (**m**), translation complex members NCBP1 (**n**) and ETF1 (**o**), and downstream effector SMG6 (**p**), between the top and bottom aneuploidy quartiles of human cancer cell lines (n=538 cell lines). Data were obtained from DepMap CRISPR screen, 22Q1 release. ****, p<0.0001 for EIF4A3, NCBP1 and ETF1 respectively, **, p=0.0041 for SMG6; two-tailed Mann-Whitney test.

**Extended Data Figure 4: Aneuploid cells activate the miRNA-mediated RNA degradation pathway, and depend on it for downregulating their gene expression (related to Fig. 3) (a)** Western blot of DROSHA protein levels in RPE1 clones treated with siRNA against DROSHA (or control siRNA) for 72hrs. GAPDH was used as a housekeeping control. **(b-c)** Comparison of gene dependency (determined by DEMETER2 score) for key members of the RISC complex, TARBP2 (**b**) and PRKRA (**c**), between the top and bottom aneuploidy quartiles of human cancer cell lines (n=252 and n=348 cell lines, for TARBP2 and PRKRA respectively). Data were obtained from the DepMap RNAi screen, 22Q1 release. ***, p=0.0007 and p=0.0004 for TARBP2 and PRKRA, respectively; Two-tailed Mann-Whitney test. **(d)** Western blot of DROSHA and γH2AX protein levels in parental RPE1 cells treated with etoposide (2.5µM) for 1, 3 or 6 hours. Elevation of DROSHA protein levels is associated with the degree of DNA damage in the cells. Tubulin was used as a housekeeping control. **(e)** Quantification of DROSHA protein levels in parental RPE1 etoposide-treated cells, calculated relative to the DMSO control per experiment. n=4 independent experiments. *, p=0.034, p=0.0375, p=0.0456 for 1h, 3h, and 6h etoposide treatment respectively; One Sample t-test. **(f)** Gene set enrichment analysis (GSEA) of splicing-related signatures, comparing the highly aneuploid clones, SS51 and SS111, to the pseudo-diploid clone SS48. Shown are enrichment plots for GO-Biological Processes ‘Regulation of splicing’ (NES=-1.925; q-value=0.109), ‘Alternative mRNA splicing via spliceosome’ (NES=-1.925; q-value=0.109), and ‘Regulation of alternative mRNA splicing via spliceosome’ (NES=-1.837; q-value=0.13) gene sets. **(g-h)** Estimation of alternative splicing activity in the near-diploid and aneuploid clones. Shown are percent spliced in (PSI) values for 3’ (**g**) and 5’ (**h**) splice site recognition (Alt3 and Alt5, respectively), obtained by applying Vast-Tool^64^ to the RNAseq data. High PSI values represent retention of splicing events in samples, thus decreasing splicing activity. Comparison of common splicing events is performed for each sample against SS48, separately. ****, p<0.0001; two-tailed Wilcoxon test.

**Extended Data Figure 5: Multiple models of aneuploid cells experience proteotoxic stress and attenuate protein translation (related to Figure 4) (a)** Comparison of drug sensitivity (determined by EC50 values) to 48hr drug treatment with the UPR activator tunicamycin, between parental RPE1 cells, and their highly-aneuploid derivatives RPTs. n=4 independent experiments. EC50 fold-change was calculated relative to RPE1, per experiment. *, p=0.0126, **, p=0.0072 and p=0.0095 for RPT3, RPT1 and RPT4, respectively; One-Sample t-test. **(b)** Representative image of SUnSET puromycin incorporation assay, showing reduction in global translation in the highly-aneuploid RPT clones in comparison to parental RPE1 cells. GAPDH was used as housekeeping control. **(c)** Quantitative comparison of SUnSET puromycin incorporation in parental RPE1 cells and derivative highly-aneuploid RPT, calculated relative to the parental cells. n=5 independent experiments; *, p=0.0115 and p=0.0149, for RPT1 and RPT4 respectively, **, p=0.0073 for RPT3; One-Sample t-test **(d)** Comparison of drug sensitivity (determined by EC50 values) to 48hr drug treatment with the UPR activator tunicamycin, between parental RPE1 cells treated for 20hrs with the SAC inhibitor reversine (500nM) or with control DMSO. n=5 independent experiments. EC50 fold-change was calculated relative to RPE1-DMSO cells, per experiment. **, p=0.0017; One-Sample t-test **(e)** Representative image of SUnSET puromycin incorporation assay between parental RPE1 cells treated for 20hrs with the SAC inhibitor reversine (500nM) or with control DMSO, showing reduction in global translation following reversine-mediated aneuploidization. Vinculin was used as a housekeeping control. **(f)** Quantitative comparison of SUnSET puromycin incorporation between DMSO and reversine-treated RPE1 cells, calculated relative to DMSO-treated cells. n=6 independent experiments; **, p=0.0012; One-Sample t-test.

**Extended Data Figure 6: Multiple models of aneuploid cells activate the proteasome, and depend on its activity for downregulating their protein expression (related to Figure 5) (a)** Comparison of mRNA levels, quantified by qRT-PCR, in parental RPE1 cells and derivative highly aneuploid RPTs of representative subunits of the 20S and 19S proteasome complexes (left to right): PSMA1, PSMB5, PSMC1, and PSMD12. Fold change in expression was calculated relative to parental RPE1, per experiment. n=6 independent experiments; *, p<0.05, **, p<0.01, ***, p<0.001; One-Sample t-test. **(b)** Comparison of mRNA levels, quantified by qRT-PCR, of representative subunits of the 20S and 19S proteasome complexes (left to right): PSMA1, PSMB5, PSMC1, and PSMD12, between parental RPE1 cells treated for 20hrs with the SAC inhibitor reversine (500nM) or with control DMSO. Fold change in expression was calculated relative to DMSO-treated cells, per experiment. n=6 independent experiments; *, p<0.05, **, p<0.01, ***, p<0.001 in each panels; One-Sample t-test. **(c)** The levels of proteasome activity, measured by Proteasome-Glo®, in parental RPE1 cells and derivative highly-aneuploid RPT cells, showing increased proteasome activity in RPT cells. Proteasome activity was calculated relative to RPE1, per experiment. n=4 independent experiment, *, p=0.022 and p=0.0264, **, p=0.006, for RPT1, RPT3, and RPT4, respectively; One-Sample t-test. **(d)** The levels of proteasome activity, measured by Proteasome-Glo®, in DMSO vs. reversine-treated RPE1 cells, showing increased proteasome activity following reversine treatment. Proteasome activity was calculated relative to DMSO-treated cells, per experiment. n=4 independent experiment; *, p=0.0342; One-Sample t-test. **(e)** Comparison of dependency scores for all proteasome subunits in aneuploid clones (SS6, SS119, SS51) vs the pseudo-diploid clone SS48. Aneuploid cells are more dependent on the proteasome subunits in comparison to pseudo-diploid clones. n=38 genes; ****, p<0.0001; two-tailed Mann-Whitney test. **(f-g)** Representative images of CAL51, a near-euploid breast cancer cell line, vs. MDA-MB-468, a highly-aneuploid breast cancer cell line, following exposure to bortezomib for 72hrs. The highly-aneuploid cell line is more sensitive to the treatment.

## Author contributions

U.B.-D. and S.S. jointly conceived the study, directed and supervised it. J.Z. and M.R.I. jointly designed and performed most of the experiments. J.Z., M.R.I., U.B.-D. and S.S. analyzed the data with inputs from all co-authors. Y.E. and E.R. assisted with bioinformatic analyses. S.V., G.D.F., A.S.K., I.V., S. M, K.L., Y. C.-S. and S.S. assisted with *in vitro* experiments. F.V. directed the genomic profiling and CRISPR screens. J.Z., M.R.I., U.B.-D. and S.S. wrote the manuscript with inputs from all co-authors.

## Materials availability

Aneuploid RPE1-hTERT clones generated in this study are available upon request to Stefano Santaguida. Raw RNAseq data are available in the SRA database (https://www.ncbi.nlm.nih.gov/sra) under accession number PRJNA889550. Genome-wide CRISPR/Cas9 screening data of RPE1-hTERT clones are available in supplementary table in Zerbib, Ippolito et al, *bioRxiv* 2023 and in the DepMap database 21Q3 release (https://figshare.com/articles/dataset/DepMap_21Q3_Public/15160110). Cancer cell line expression, CRISPR/Cas9 and RNAi data are available in the DepMap database 22Q1 release (https://figshare.com/articles/dataset/DepMap_22Q1_Public/19139906). All of them are publicly available as of the date of publication. Aneuploidy scores of cancer cell lines are available in Zerbib, Ippolito et al, *bioRxiv* 2023.

## Supplementary information

**Supplementary Table 1: PRISM screen results of human cancer cell lines treated with bortezomib, in the absence or presence of reversine** Comparison of the EC50 values of bortezomib in human cancer cell lines in the absence or presence of a sub-lethal dose (250nM) of reversine (or vehicle control) for 5 days.

**Supplementary Table 2: Details of reagents and oligonucleotides used in the study**

