## Supplementary figures and images for "Increased RNA and protein degradation is required for counteracting transcriptional burden and proteotoxic stress in human aneuploid cells"

### Extended Data Fig. 1

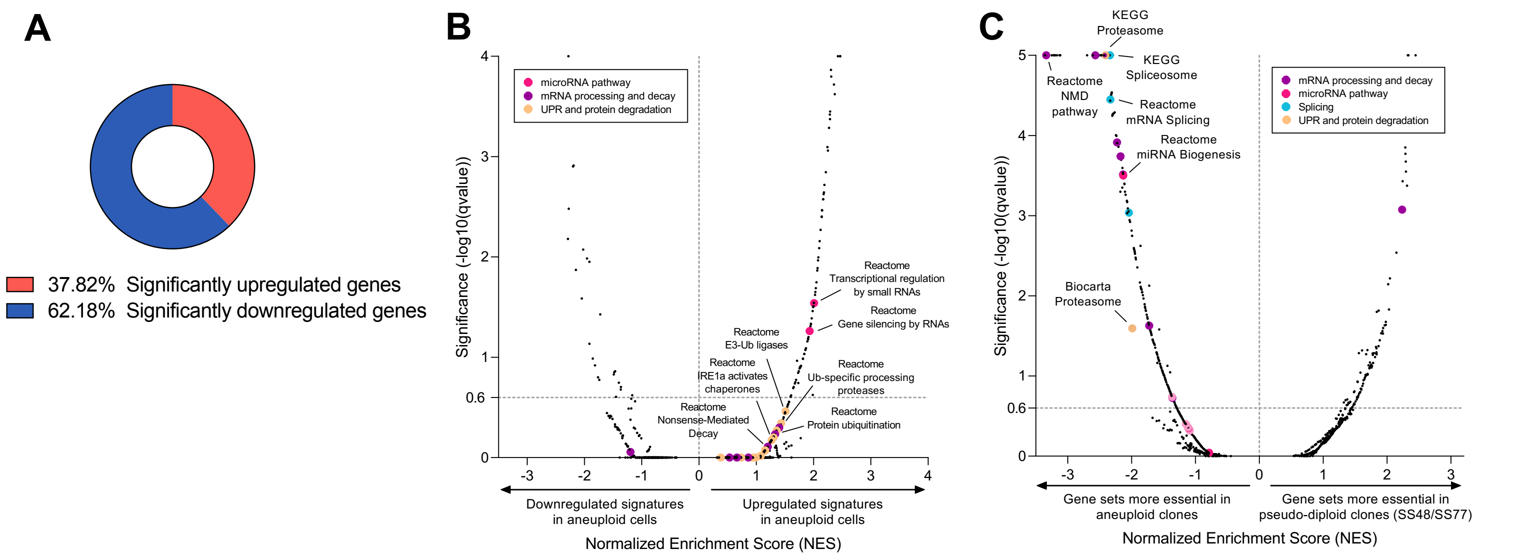


**Extended Data Figure 1**

### Extended Data Fig. 2

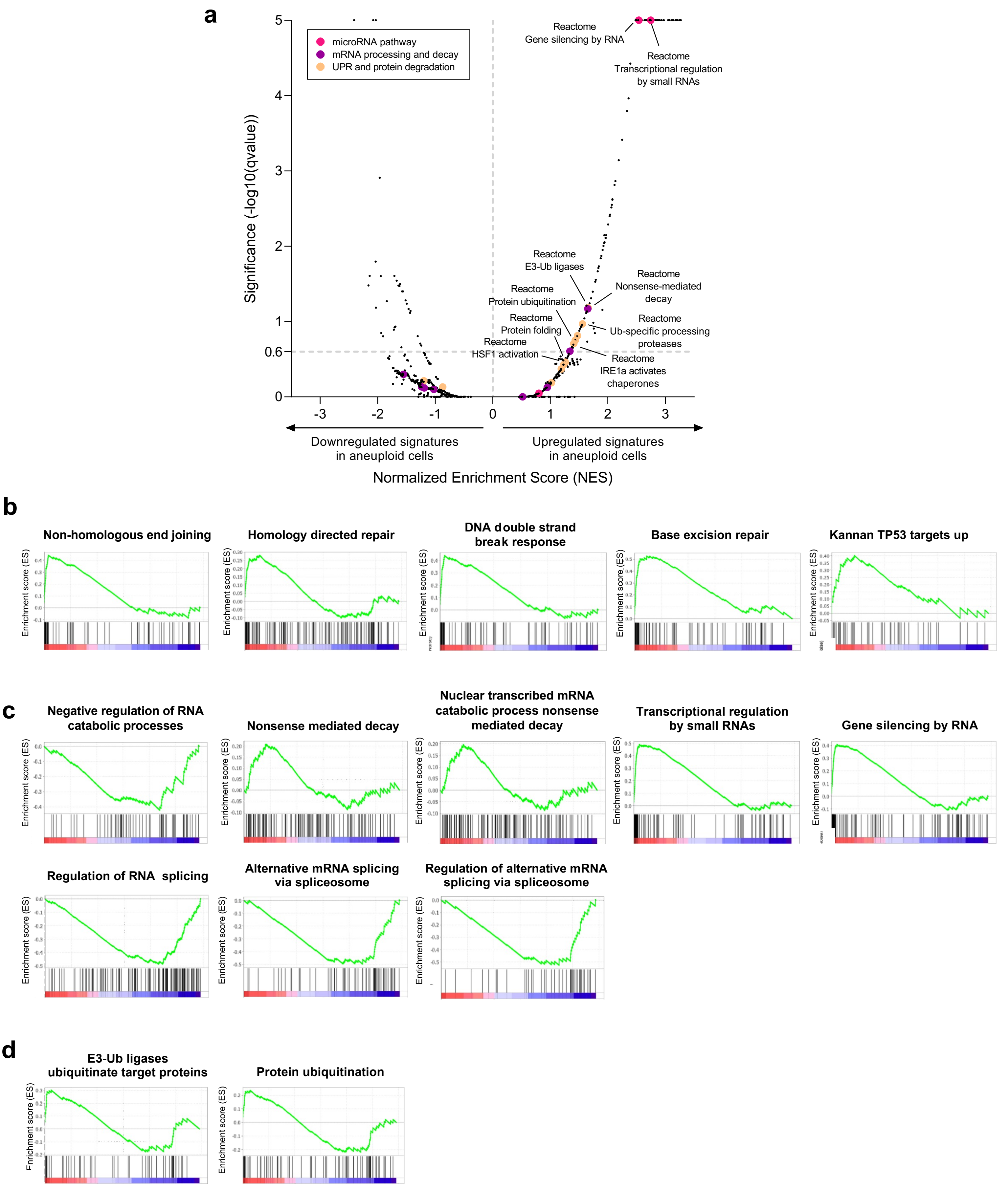


**Extended Data Figure 2**

### Extended Data Fig. 3

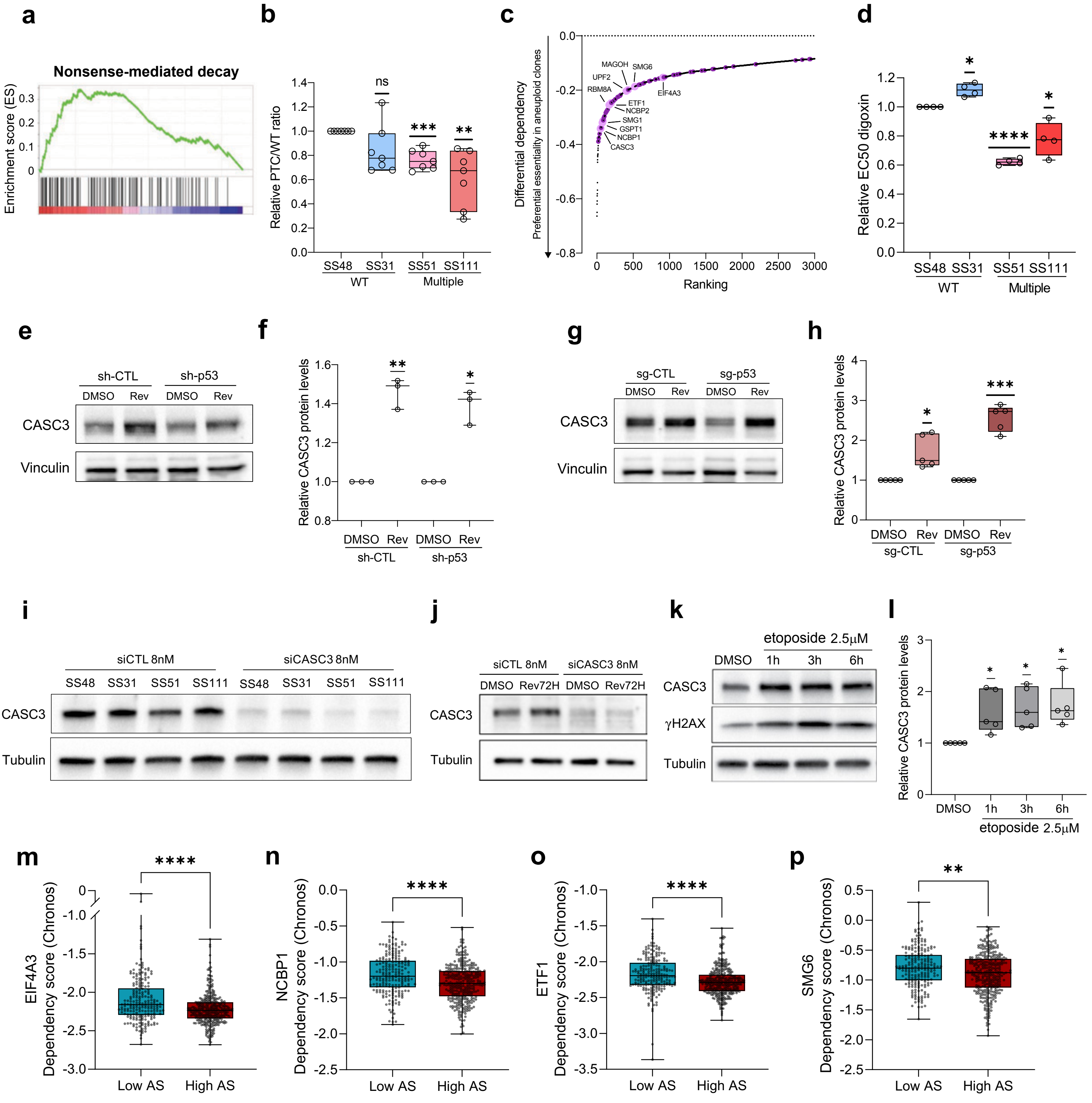


##### Extended Data Figure 3

### Extended Data Fig. 4

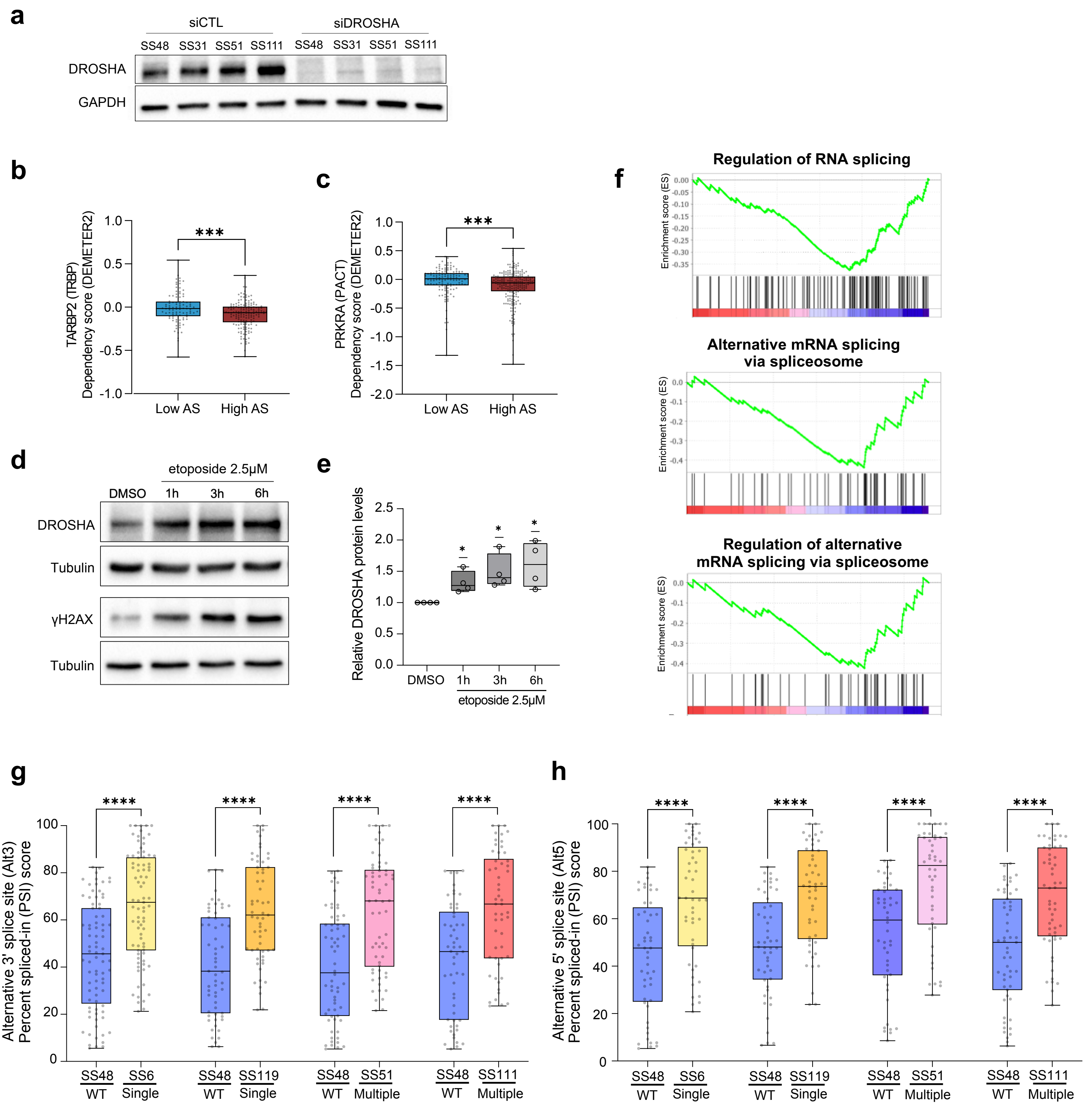


Extended Data Figure 4

### Extended Data Fig. 5

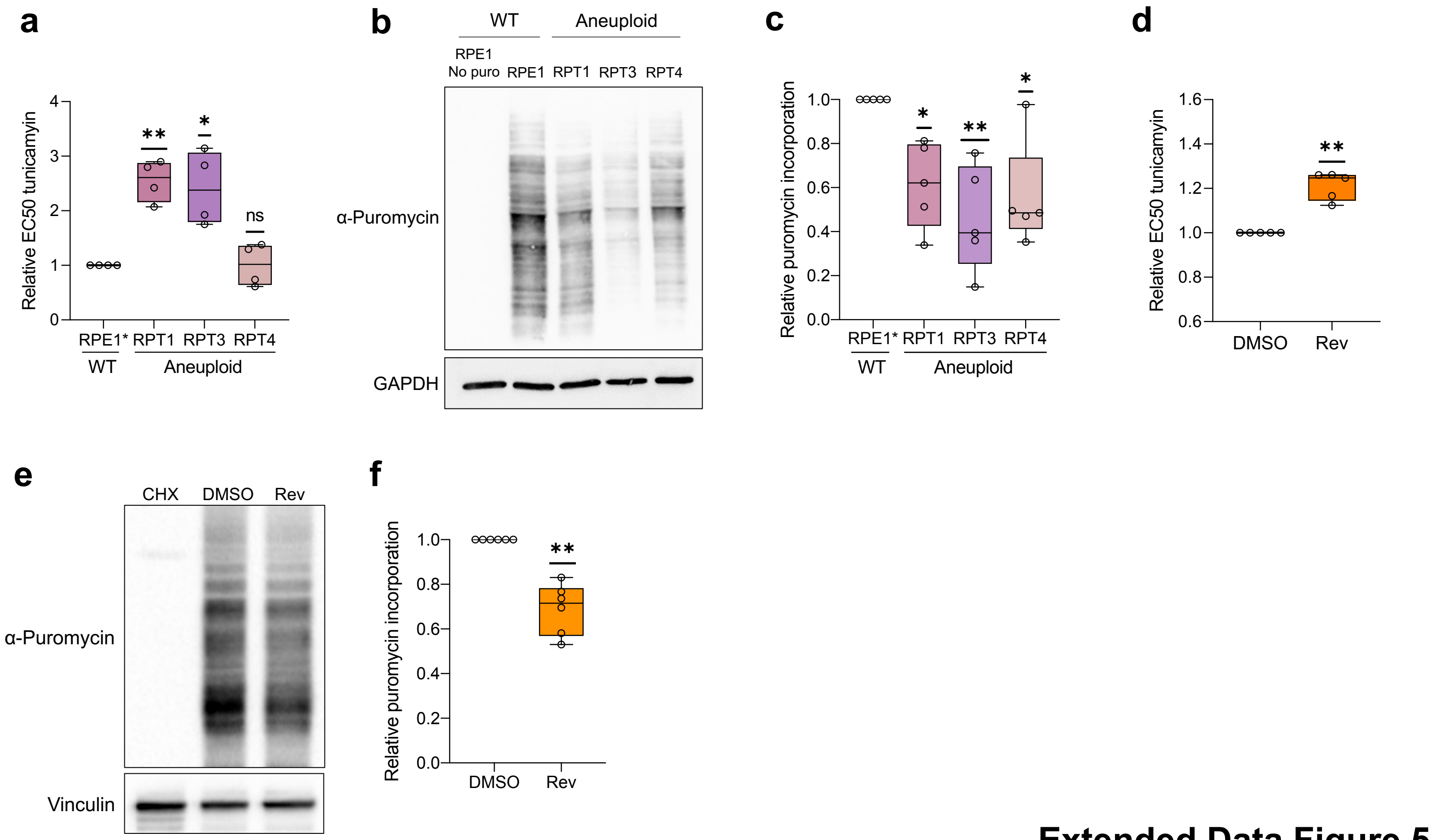


**Extended Data Figure 5**

### Extended Data Fig. 6

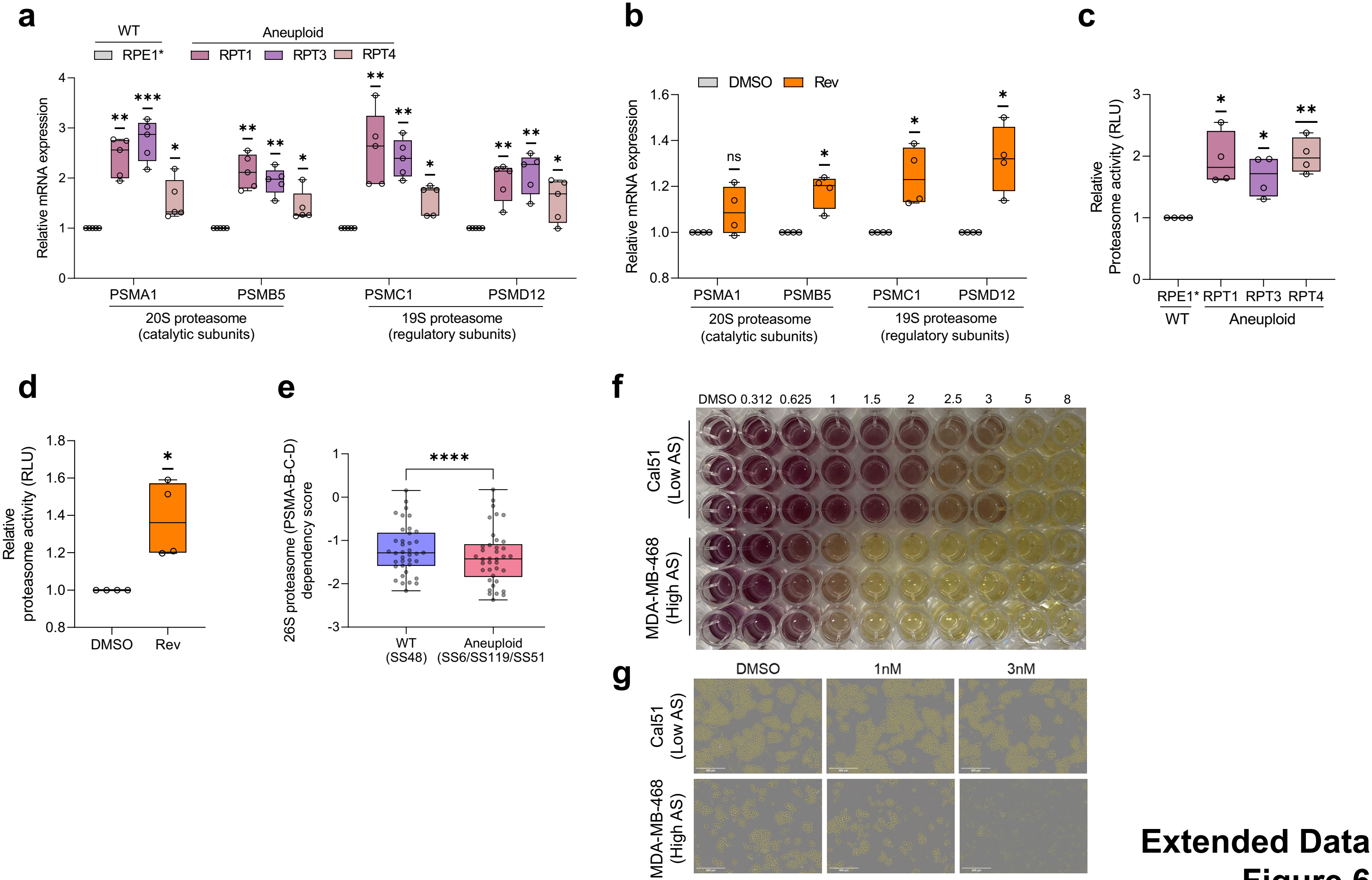


#### Extended Data Figure 6
